# Common DNA sequence variation influences epigenetic aging in African populations

**DOI:** 10.1101/2024.08.26.608843

**Authors:** Gillian L. Meeks, Brooke Scelza, Hana M. Asnake, Sean Prall, Etienne Patin, Alain Froment, Maud Fagny, Lluis Quintana-Murci, Brenna M. Henn, Shyamalika Gopalan

**Affiliations:** Integrative Genetics and Genomics Graduate Program, University of California, Davis, CA 95694, USA; Department of Anthropology, University of California, Los Angeles, CA, 90095, USA; Forensic Science Graduate Program, University of California, Davis, CA, 95694, USA; Human Evolutionary Genetics Unit, CNRS UMR2000, Paris, 75015, France; Institut de Recherche pour le Développement, UMR 208, Muséum National d’Histoire Naturelle, Paris, 75005, France; Université Paris-Saclay, INRAE, CNRS, AgroParisTech, Genetique Quantitative et Evolution - Le Moulon, Gif-sur-Yvette, 91190, France; Chair Human Genomics and Evolution, Collège de France, Paris, 75231, France; Department of Anthropology, University of California Davis, Davis, CA, 95616, USA; UC Davis Genome Center and Center for Population Biology, University of California, Davis, CA 95694, USA; Department of Ecology and Evolution, Stony Brook University, Stony Brook, NY, 11790, USA; Department of Genetics and Biochemistry, Clemson University, Clemson, SC 29634, USA; Center for Human Genetics, Clemson University, Greenwood, SC 29646, USA

## Abstract

Aging is associated with genome-wide changes in DNA methylation in humans, facilitating the development of epigenetic age prediction models. However, most of these models have been trained primarily on European-ancestry individuals, and none account for the impact of methylation quantitative trait loci (meQTL). To address these gaps, we analyzed the relationships between age, genotype, and CpG methylation in 3 understudied populations: central African Baka (n = 35), southern African ‡Khomani San (n = 52), and southern African Himba (n = 51). We find that published prediction methods yield higher mean errors in these cohorts compared to European-ancestry individuals, and find that unaccounted-for DNA sequence variation may be a significant factor underlying this loss of accuracy. We leverage information about the associations between DNA genotype and CpG methylation to develop an age predictor that is minimally influenced by meQTL, and show that this model remains accurate across a broad range of genetic backgrounds. Intriguingly, we also find that the older individuals and those exhibiting relatively lower epigenetic age acceleration in our cohorts tend to carry more epigenetic age-reducing genetic variants, suggesting a novel mechanism by which heritable factors can influence longevity.

## Introduction

The aging process is associated with significant, genome-wide epigenetic changes. In particular, DNA methylation levels at specific cytosine-guanine dinucleotides (CpGs) have been shown to be strongly associated with chronological age, driving the development of a suite of age prediction algorithms referred to as ‘epigenetic clocks’. While thousands of CpG sites across the genome exhibit consistent patterns of increasing or decreasing DNA methylation with age^2–4^, accurate age predictors have been constructed from remarkably few CpGs^5–12^. The first DNA methylation-based predictors were trained on individuals’ chronological age (i.e. the actual number of years lived), and found that epigenetic clocks could be more accurate and precise than other molecular methods of age estimation, such as telomere length^13–15^. Interestingly, subsequent research found that the error in epigenetic clock-based age estimates (i.e. the deviation between true and predicted age) is also biologically meaningful, and that an accelerated epigenetic age is associated with multiple age-related diseases^16–19^. This observation spurred the next generation of epigenetic predictors, which included PhenoAge^20^, GrimAge^21,22^, and FitAge^23^, that were specifically trained to predict morbidity, mortality, and other aspects of biological aging.

Deviation between one’s predicted and actual age, i.e., epigenetic age acceleration, has been shown to be influenced by a host of environmental and lifestyle factors^24^, leading researchers to examine its relationship to systemic health disparities experienced by minorities in cosmopolitan populations^25–32^. However, these epigenetic models are almost exclusively trained on European-descent populations living in industrialized societies, and are rarely validated across a range of genetic backgrounds and environmental contexts. In fact, studies that have assessed popular predictors in genetically diverse cohorts often find inconsistent patterns. For example, both African-American and Hispanic cohorts exhibit systematically higher epigenetic age under some epigenetic models, and systematically lower epigenetic age under others^20,32–34^ ^20,33–35^. While these inconsistencies might reflect real among-population variation in the aging process, without first confirming that epigenetic clocks maintain their predictive power across diverse human populations, researchers should be cautious about interpreting the causes and consequences of epigenetic age acceleration in relation to human health^36^.

A recent study found that prediction accuracy does indeed decline when clocks trained on a particular genetic ancestry are applied to individuals of a diverged genetic ancestry^37–39^, mirroring findings from studies of polygenic risk score (PRS) transferability^40,10,37–39,41^. This observation might be partially explained by the relatively high heritability of DNA methylation and the strong influence of individual single nucleotide polymorphisms (SNPs)^42–44^; 10% of CpG sites exhibit heritability greater than 50%, and up to 45% of CpG sites assayed by the Illumina 450k array show influence of methylation quantitative trait loci (meQTL), 90% of which act locally, in *cis*^45^. This significant genetic control of DNA methylation also explains why genome-wide variation in DNA methylation broadly recapitulates patterns of population structure observed in human genetic data^46–51^.

Previous work has identified meQTL as being important drivers of variation at some age-associated CpGs^13,14,52^. If these same CpGs are used to construct an epigenetic clock, its accuracy might be expected to decline when applied to a genetically diverged population. This is because predictor coefficients will be biased based on the meQTL frequency of the training cohort.

In order to address these questions, we assessed several popular epigenetic clocks on genetically diverse populations, and characterized the influence of genetic variation on DNA methylation both within and across populations. We analyze saliva-derived DNA methylation data from three African populations representing a broad swath of different genetic ancestries: Baka central African foragers, southern African ‡Khomani San foragers, and southern African Himba pastoralists. Each of these groups have a distinct, complex evolutionary history and currently occupy very different ecological regions across the continent, generating among-population variation in both genetic and environmental factors that can influence DNA methylation. We compare the predictive accuracy of 10 published epigenetic clocks on these African cohorts, as well as on publicly available data from European-ancestry and Hispanic/Latino cohorts^35^. Using paired genotype data for the African individuals in our dataset and newly-available, ancestry-matched imputation panels^53^, we estimated heritability and identified significant *cis*-meQTL for age-associated CpGs across the genome. Importantly, we find that a large proportion of CpGs included in established predictors are influenced by meQTL identified in our modestly-sized cohorts. We show that not accounting for genetic variation at meQTL contributes to error in epigenetic age prediction, and develop novel epigenetic clocks that specifically exclude CpGs with significant *cis*-heritability. Finally, we develop a genotype-based ‘epigenetic aging score’ (EAS), which captures the effects of epigenetic age-increasing DNA variants from across the genome under an additive model and find correlations with independently derived estimates of epigenetic age acceleration, suggesting biologically meaningful effects at some of these meQTL (Supplementary Figure 1).

## Results

### Evaluating performance of published age predictors on African cohorts

We tested 10 age prediction methods (see Methods) that were trained primarily on European-ancestry cohorts living in Europe and the United States, the Horvath multi-tissue age predictor^13^, the Hannum blood clock^14^, the Horvath skin and blood clock^54^, the Zhang elastic net predictor^55^, PhenoAge^20^, two iterations of GrimAge both using either true or predicted age^21,22^, and FitAge^23^. We applied the predictors to saliva-derived DNA methylation data from 3 African cohorts and compared performance to a publicly available tissue-matched dataset of European-ancestry and Hispanic/Latino individuals (GEO accession GSE78874^35^). Because some clocks show age-dependent accuracy^56^, we primarily report age-adjusted prediction errors to account for the different age distributions across cohorts. We found that 9 of the predictors exhibited significant differences in age-adjusted error between at least one African population and the European and Hispanic/Latino datasets (Figure 1; Supplementary Tables 1-2; Supplementary Figure 2). There was not a consistent pattern of over- or under-estimation for the African cohorts relative to the European and Hispanic/Latino individuals; for example, the Himba as a group were estimated to be younger than Europeans by most clocks, but older by GrimAge2 based on true age; ‡Khomani San individuals were estimated to be younger than Europeans by the Hannum and Zhang clocks, but older by FitAge and GrimAge. We also found significant differences in prediction error among the three African cohorts. Only the Horvath clock showed no differences in age-adjusted error in the African samples as compared to the European and Hispanic/Latino samples.

**Figure 1.**
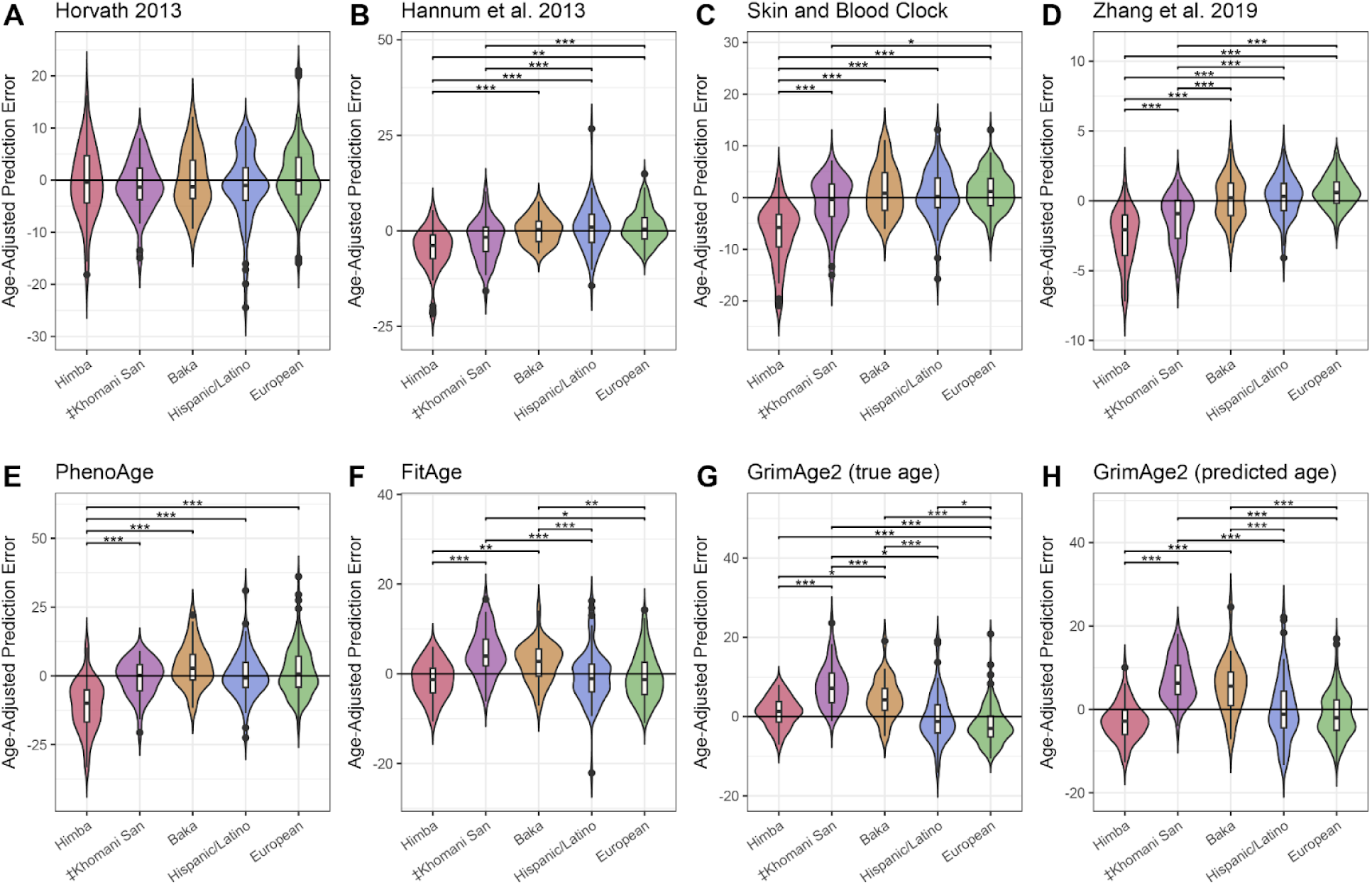
Distributions of age-adjusted prediction error across diverse cohorts. Violin plots A-H show differences in prediction error, adjusted for individual age, among Himba, ‡Khomani San, Baka, European, and Hispanic/Latino samples across 8 published epigenetic clocks. We tested for significant differences in age-adjusted prediction error among all populations by ANOVA, followed by a Tukey test to identify significant pairwise differences. * indicates an adjusted p-value of < .05, ** < .01, and *** < .001.

We considered that differences in predictive accuracy might be due to cryptic differences in cell-type composition. Although all the samples were nominally saliva-derived and we restricted comparisons to samples predicted to be saliva or blood-derived (see Methods), significant among-population variation in the proportions of white blood cell types and epithelial cells might still exist. With the exception of the Horvath multi-tissue clock^13^ and the skin and blood clock^54^, the predictors that we evaluated were trained primarily on whole blood-derived DNA methylation data, and are not expected to perform uniformly well across tissues. Therefore, if the cell-type composition of samples varied systematically across cohorts, this could produce differences in predictive accuracy that appear to be population specific. We were especially concerned that the high frequency of the Duffy null variant in West African populations^57^, which is associated with lower neutrophil count in whole blood^58–60^, could also drive ancestry-associated differences in saliva cell-type composition.

As expected^61^, we found the Duffy null variant is fixed, or nearly fixed, in the Himba and Baka (allele frequency of 100% and 94%, respectively). The frequency of Duffy null in the ‡Khomani San cohort was 27%, consistent with gene flow from West African ancestry populations into an environment where selection for malarial resistance is low^62^. Because of this intermediate frequency, we were able to test for a relationship between Duffy null genotype and estimated neutrophil proportion, as well as with overall predictive accuracy, within the ‡Khomani San cohort. We applied a reference-based cell-type deconvolution method^63^ to estimate cell-type proportions in each sample (see Methods)^64^. We observed a slight, but non-significant negative relationship between neutrophil proportion and Duffy genotype in the ‡Khomani San and Baka cohorts (Supplementary Figure 3). However, we did not find that this led to a significant difference in prediction error across the 10 predictors (Supplementary Figure 4).

Interestingly, there were fewer significant pairwise differences among cohorts across 10 different measures of epigenetic age acceleration (Supplementary Figure 5). Most of these measures were derived from the Horvath^13^, Hannum^14^, PhenoAge^20^, and GrimAge^21,22^ clocks, while one was developed independently as a DNA methylation-based estimate of the rate of telomere shortening^65^ (see Methods). Based on the PhenoAge and GrimAge-based epigenetic age acceleration metrics, the Himba and Baka had significantly higher acceleration than the European samples (Supplementary Figure 5). In line with this, the Himba and Baka also had significantly shorter methylation-based estimates of telomere length for their age than the European and Hispanic cohorts.

### Epigenome-wide association study for age

We conducted an epigenome-wide association study (EWAS) to identify CpG sites whose methylation levels are associated with age for each of our three population cohorts. We identified 347, 149, and 282 CpG sites that met the Bonferroni-corrected threshold for significance in the Himba, ‡Khomani San, and Baka, respectively for a total of 657 unique sites. 31 of these sites were identified independently in all three populations. We found that the estimated effect sizes were highly correlated in all three pairwise comparisons after conditioning on significance in at least 1 of the populations (Figure 2A-C).

**Figure 2.**
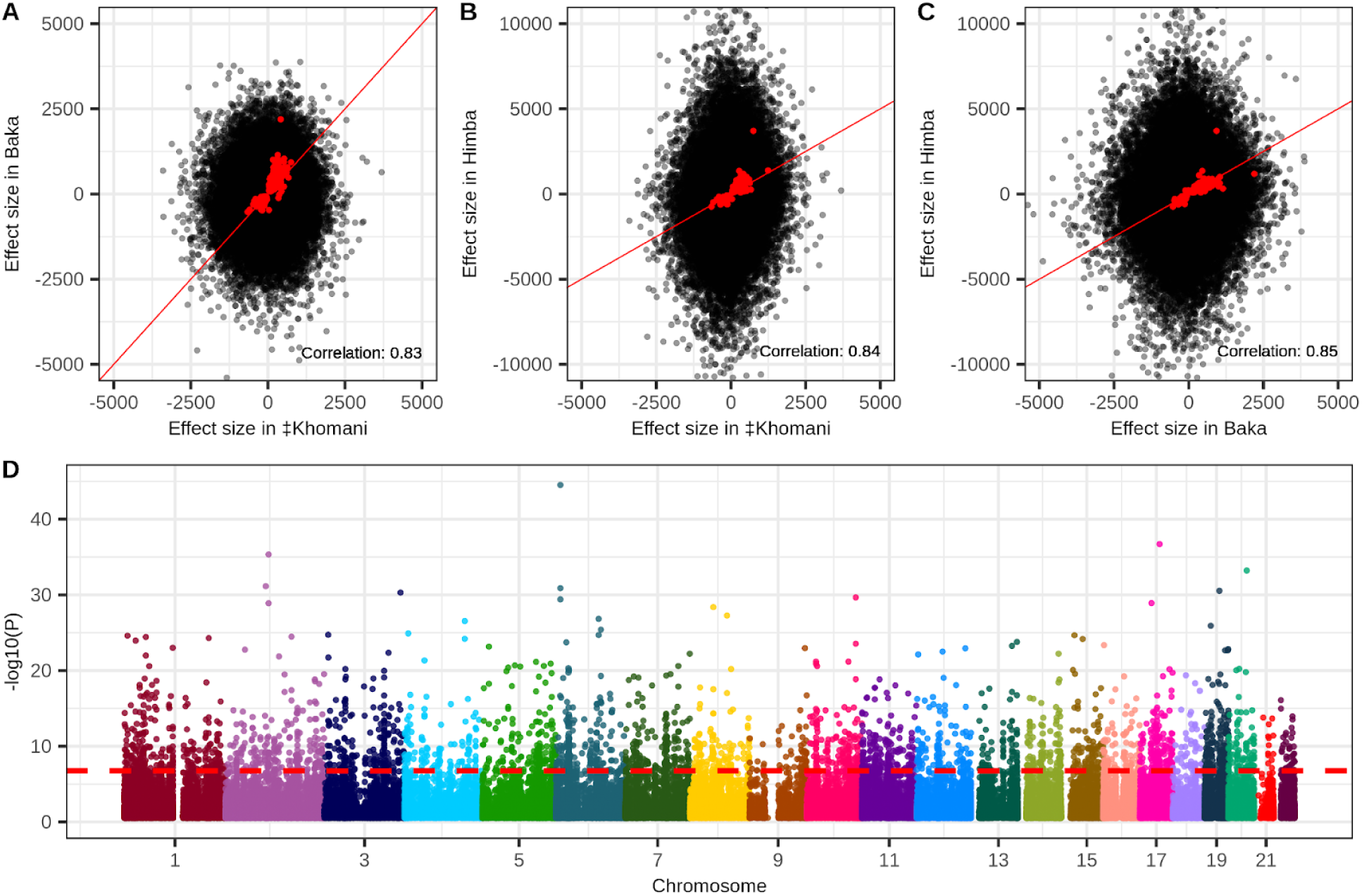
Epigenome-wide associations with chronological age. Panels A-C show the correlation of estimated effect sizes of DNA methylation on age for all pairwise comparisons of the Baka, ‡Khomani San, and Himba. Red points indicate CpG sites that were identified as significantly associated with age in at least one of the three populations. For these points the correlation between effect sizes estimated in different populations is indicated at the bottom right of each panel. Panel D is a Manhattan plot depicting the strength of association with age along the entire genome from a meta-analysis of the individual epigenome-wide association studies run in the three populations. A total of 3,211 CpG sites exceed the threshold for significance.

We next conducted a fixed effect meta-analysis of our three populations to maximize our power to detect DNA methylation-age associations. Our meta-analysis identified 3,211 significant age associations across the 355,103 CpG sites common to all three datasets (Figure 2D). We found that 1,637 of these overlapped with previously identified age-associated CpG sites identified in 35 published studies (Supplementary Table 3), including our previous study which included the Baka and ‡Khomani San datasets^66^.

### Identification of cis-meQTL associations in African data

Next, we identified meQTL that influence DNA methylation in our cohorts in order to understand the impact of nearby heritable variation on age-associated CpG sites. We conducted a ‘baseline’ *cis*-meQTL scan of each African cohort separately by testing a set of common, LD-pruned variants falling within 200kb of each CpG site for association with DNA methylation level (see Methods). We identified 75,120, 61,525, and 198,775 significant meQTL in the ‡Khomani San, Baka, and Himba, respectively, affecting 11.1% (46,441), 8% (32,167), and 11.7% (83,527) of assayed CpG sites (Supplementary Figure 6B). We then assessed the overlap of CpGs influenced by *cis*-meQTL and those whose DNA methylation levels are associated with age in our meta-analysis EWAS results. We found that 645 of the 3,211 (20.1%) significant sites from the meta-analysis are influenced by an meQTL identified in at least one population. Because our starting SNP sets were different for each population and were LD pruned independently, the same SNP was rarely identified across multiple populations; however, we identified thousands of CpG sites that were influenced by meQTL in at least two populations (Supplementary Figure 6B). In cases where the same SNP was identified as a significant meQTL we found that their effect sizes were very highly correlated across populations (Figure 3A).

**Figure 3.**
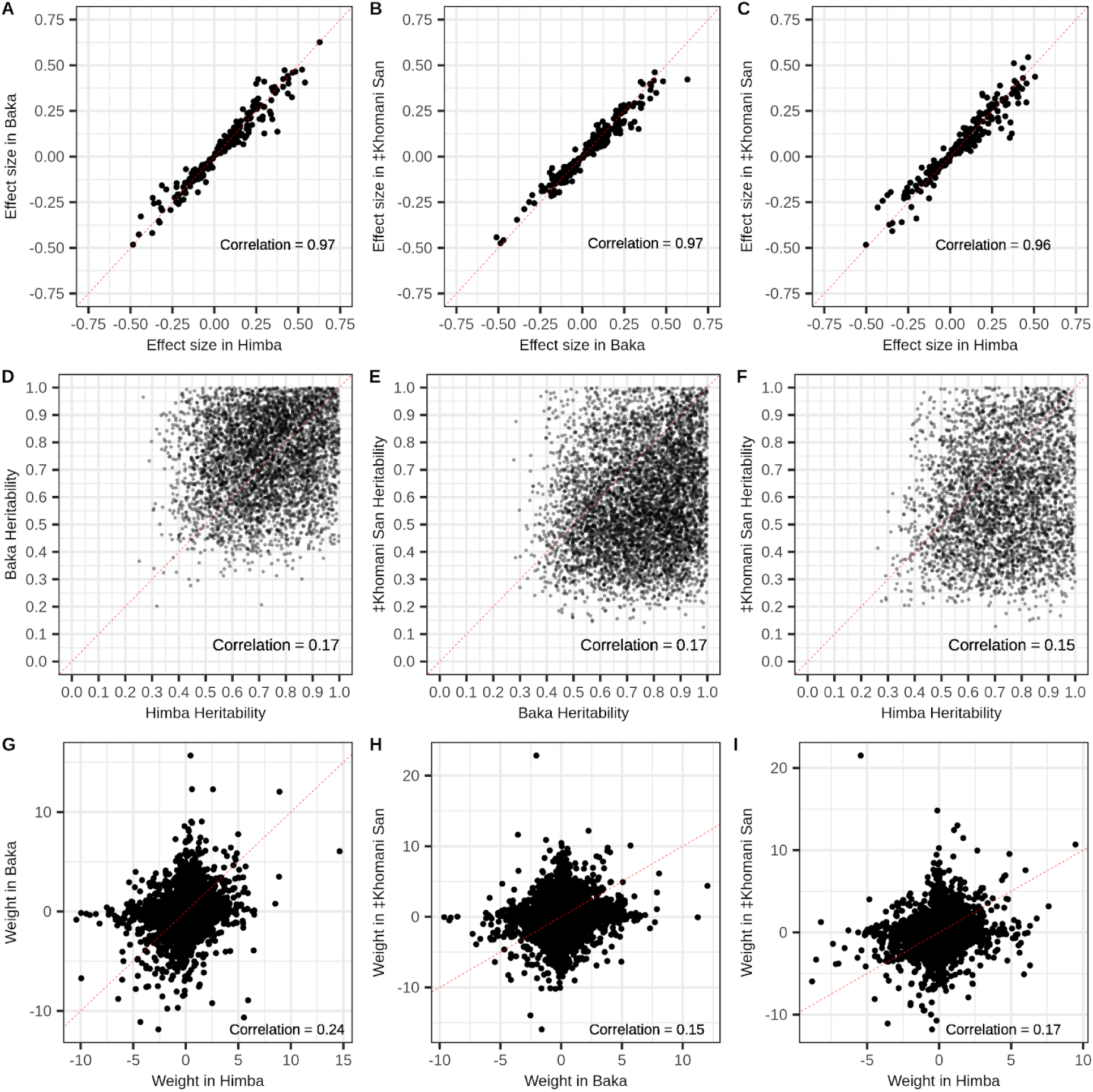
Shared *cis*-genetic architecture of CpG methylation among populations. Panels A-C show the correlations of estimated effect sizes of SNP genotype on DNA methylation level from baseline *cis*-meQTL scans of the Baka, ‡Khomani San, and Himba for cases where the same SNP-CpG relationship was identified in both populations. Panels D-F show the correlation in *cis*-heritability measures for significantly heritable (p value < .05) CpG sites across all pairwise combinations of populations. Panels G-I show the correlations of *cis*-SNP weights on DNA methylation levels estimated from the FUSION regression models for the instances where the same SNP was estimated to have a non-zero weight across different populations, but the selected model was allowed to vary between populations. Weights were scaled within model typeIn each panel, the dashed represents the line of equality.

We also conducted an ‘extended’ *cis*-meQTL analysis using the FUSION^67^ software package that considered a 1Mb window around each CpG site to first estimate *cis*-heritability and then for significantly heritable sites, model SNP weights using 4 regression methods: elastic net, LASSO^68^, SuSie (sum of single-effect)^69^, and best single meQTL (see Methods)^67^. We used non-LD pruned genotype data for this analysis to gauge the extent to which the genetic architecture of CpG methylation is conserved across populations. We then moved forward with the best performing of the 4 regression models for each individual CpG site (Supplementary Table 4).

Even with our modest sample sizes, we found that a substantial proportion of CpG sites (8.2%, 6.7%, and 10.2% of CpGs tested in the ‡Khomani San, Himba, and Baka, respectively) exhibited significant *cis*-heritability (p < 0.05); 8.2%, 6.7%, and 10.2% of CpG sites tested in the ‡Khomani San, Himba, and Baka, respectively, were heritable (Supplementary Figure 6C). Furthermore, for significantly heritable sites *cis-*heritability of CpG methylation was significantly, but weakly correlated across all pairs of populations (Pearson correlations: Himba-Baka r = .17; Baka-‡Khomani San r = .17; Himba-‡Khomani San r = .15) (Figure 3B). We also tested the correlation of non-zero SNP weights across population pairs, scaling weights within each regression model type, when the same SNP was reported to have a non-zero weight in multiple populations. We expected correlations to be lower than in the baseline meQTL scan as different regression models could be selected as the best performing model across populations. Weights determined by the models allowing for multiple SNP effects (elastic net, LASSO, and SuSie) are also dependent on the specific cis-variant sets in each population and would lead to lower correlations of weights across populations. We did, however, still find moderately correlated effect sizes (Pearson correlations: Himba-Baka r = .24 ; Baka-‡Khomani San r = .15 ; Himba-‡Khomani San r = .17) (Figure 3C).

### Accounting for *cis*-genetic influence in EWAS improves associations

Since meQTL variation is generally expected to add noise to the relationship between CpG methylation and age^14,66^, we reasoned that regressing out SNP effects for significant meQTL from the corresponding DNA methylation values should tend to improve age associations. To test this, we re-ran our population-specific EWAS after first regressing out the effect of genotype at the top meQTL identified by our baseline scan from its respective CpG site’s DNA methylation values. As expected, this approach resulted in a greater number of CpG sites passing the significance threshold compared to the original EWAS: 164 (increase of 15), 405 (increase of 58), and 312 (increase of 30), in the ‡Khomani San, Himba, and Baka respectively (Supplementary Figure 7A-F). This represents 751 unique sites identified across the three populations, an additional 96 sites compared to the original EWAS. 655 of the 657 original unique associations were replicated in at least one population in the meQTL-regressed EWAS.

We conducted a meta-analysis on the meQTL-regressed EWAS results and found 3,427 significant associations, including 224 CpG sites that were not significant in the original meta-EWAS (Supplementary Figure 7G-H). 3203 of the 3211 associations identified in the initial meta-EWAS remained significant in the meQTL-regressed meta-EWAS. We found that for the 645 CpG sites that were significantly associated with age in the original meta EWAS and also influenced by meQTL, 34.3% showed an improved association with age by a p-value reduction of at least one order of magnitude.

Across all CpG sites influenced by meQTL, 2.6%, 4.7%, 6.8%, and 2.4% improved their association with age by at least one order of magnitude in the ‡Khomani San, Himba, Baka, and meta-analysis, respectively (Figure 4A-D). In order to ensure that this observed improvement was not spurious, we conducted a permutation analysis where we instead regressed out genotype values for a random SNP from a different chromosome. Across 100 permutations, only .08%, .03%, .06%, and .03% of CpG sites (Figure 4A-D), on average, exhibited a similar magnitude of improvement, indicating that accounting for real meQTL associations does indeed improve our power to detect the relationship between CpG methylation and age (Figure 4E-G).

**Figure 4.**
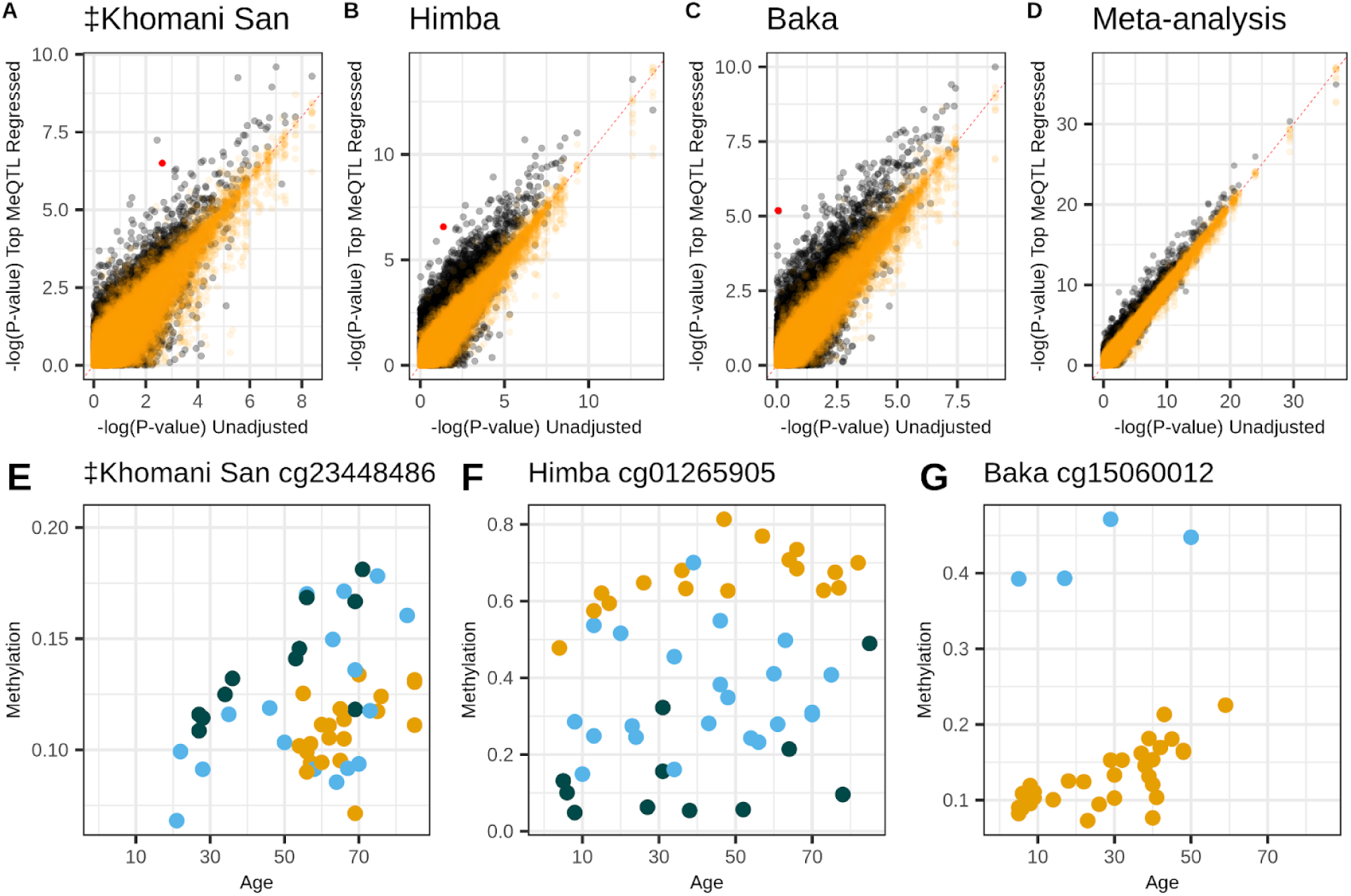
Accounting for meQTL genotype improves power to detect age associations. Panels A-D show p-values for the association between CpG methylation and chronological age from the unadjusted epigenome-wide association study (x-axes) versus p-values from the meQTL-adjusted epigenome-wide association study (y-axes). Orange points show the results of 100 permutations where a random SNP’s genotype is regressed out rather than the true meQTL. Panels D-F highlight the red points from A-C, respectively, illustrating the influence of genotype on DNA methylation at CpG sites showing particularly large p-value improvements in the adjusted EWAS. The three colors indicate the three genotype classes possible for each meQTL.

### *Cis*-meQTL influencing popular epigenetic clocks are differentiated across populations

If meQTL influence a significant proportion of CpG sites used as predictors in epigenetic clocks this will lead to increased prediction error in population samples with divergent meQTL frequencies as predictor coefficients will be calibrated based on the average meQTL genotype of the training data. This is expected to lead to particularly poor performance in out-of-sample prediction when an meQTL is very rare or invariant in the training data, but has common segregating variation in the target population. This is precisely the case in our study, as most published epigenetic clocks are trained on European-ancestry cohorts, but are being applied in African populations that have higher overall levels of heterozygosity^70^.

We assessed the proportion of CpGs included in 6 of the published predictors that are influenced by meQTL identified in our baseline and extended scans; we excluded GrimAge and GrimAge2 as the details of these models are not publically available. We found that between 22% and 43% of CpGs comprising the tested predictors are influenced by meQTL (Appendix Table 1).

We next investigated meQTL allele frequencies in the 3 African populations and European populations from the 1000 Genomes Project (Phase 3 European super-population, n = 2504 individuals). We limited this analysis to meQTL discovered from the baseline meQTL scans and aligned both 1000 Genomes and the African populations’ reference and alternate alleles to match hg37. We found that the meQTL influencing published epigenetic age predictors were often highly differentiated between European ancestry and our African populations (Figure 5A-C). On average, these meQTL had a 13%, 10.6%, and 12.4% difference in frequency in the

**Figure 5.**
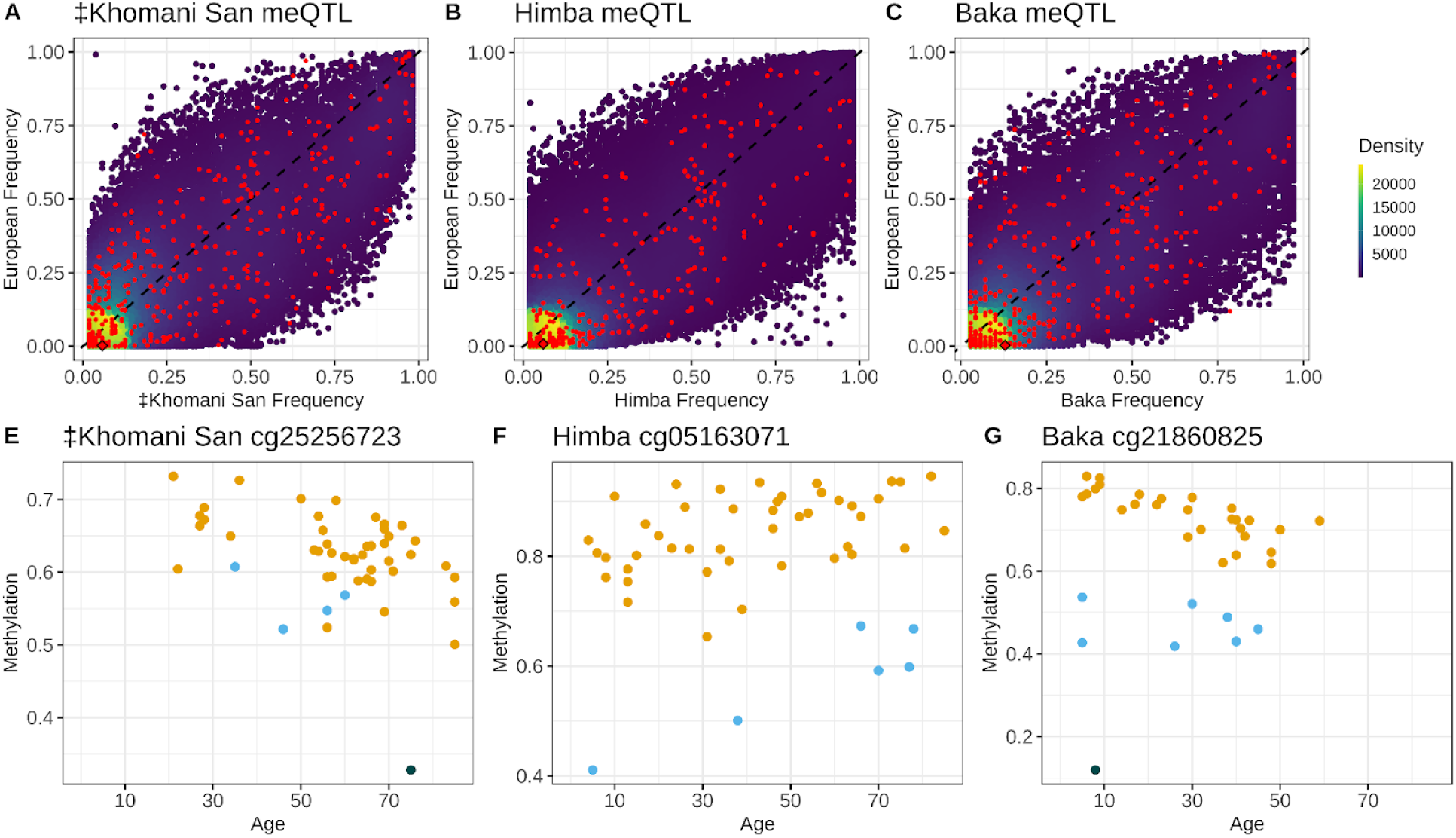
Allele frequencies of meQTL influencing CpG predictors in published epigenetic clocks are differentiated across human populations. Panels A-C show density plots of the allele frequencies of meQTL identified in each of the three African populations relative to their frequency in 1000 Genomes Phase 3 Europeans. Red points are meQTL influencing CpGs in published age prediction models. Panels D-F show the influence of genotype on baseline methylation level for the meQTL highlighted with a diamond from the top row. The three colors indicate the three possible genotype classes for each meQTL.

‡Khomani San, Himba, and Baka compared to Europeans (mean F_ST_ of .09, .11, and .12). Importantly, 5.2%, 6.7% and 9.2% of these meQTL are rare (< 1% frequency) or invariant in Europeans, but common (>5% frequency) in the ‡Khomani San, Himba, and Baka. The CpGs influenced by these meQTL would show particularly poor relative performance in non-European samples for which these variants segregate at common frequencies (Appendix Table 2). These proportions are likely an underestimate, as up to 3.5% of the meQTL we identify in the ‡Khomani San, Himba, and Baka were not present in the 1000 Genomes quality controlled, biallelic variant set and thus were not included in this analysis.

### Epigenetic clock performance is improved by excluding the effects of heritable variation

Given that variation at meQTL can reduce power to detect age associations, and that meQTL frequencies can vary substantially across human populations (Figure 5A-C), it seems prudent to exclude CpG sites under known *cis*-genetic influence when developing epigenetic clocks; this should not only optimize within-cohort performance, but also out-of-cohort transferability. In order to test this hypothesis, we used elastic net regression to construct two novel epigenetic clocks using the combined data from all three African populations to maximize our power and reduce overfitting to any one population. Of all the CpG sites common to all three populations plus the out-of cohort sample, we allowed the model to select either from CpG sites without significant *cis*-heritability, that is, not significantly influenced by a *cis*-meQTL in our baseline scans and not significantly *cis*-heritable as determined by GCTA in any of the African populations (n = 227,770), or from CpGs significantly influenced by *cis*-meQTL or determined to have significant *cis-*heritability in any of the African populations (n = 123,194). We refer to these as our “non-heritable” and “heritable” epigenetic clocks, respectively.

Supporting this hypothesis, we found that the non-heritable epigenetic clock was more accurate than the heritable epigenetic clock both in our test subset of the Himba, ‡Khomani San, and Baka, as well as in our out-of-cohort validation set of European- and Hispanic/Latino-ancestry individuals (Table 1; Figure 6). The non-heritable epigenetic clock exhibits predictive performance in African, European-ancestry, and Hispanic/Latino cohorts that is comparable to the reported test sample errors in the original Horvath and Hannum et al. publications^13,14^ (Table 1; Figure 6). Overall, these results support our hypothesis that heritable variation at meQTL negatively impacts both the transferability of epigenetic clocks as well as their overall predictive performance.

**Table 1.** Mean absolute prediction error for the non-heritable and heritable epigenetic prediction models.

|  | Non-heritable predictors | Heritable predictors | P-value* |
| --- | --- | --- | --- |
| **African test samples** | 4.27 years (.46) | 4.6 years (.52) | 5.0e-06 |
| **European samples** | 4.97 years (.87) | 5.93 years (1.64) | 7.3e-07 |
| **Hispanic/Latino samples** | 6.2 years (.98) | 6.56 years (1.27) | 2.7e-02 |

**Figure 6.**
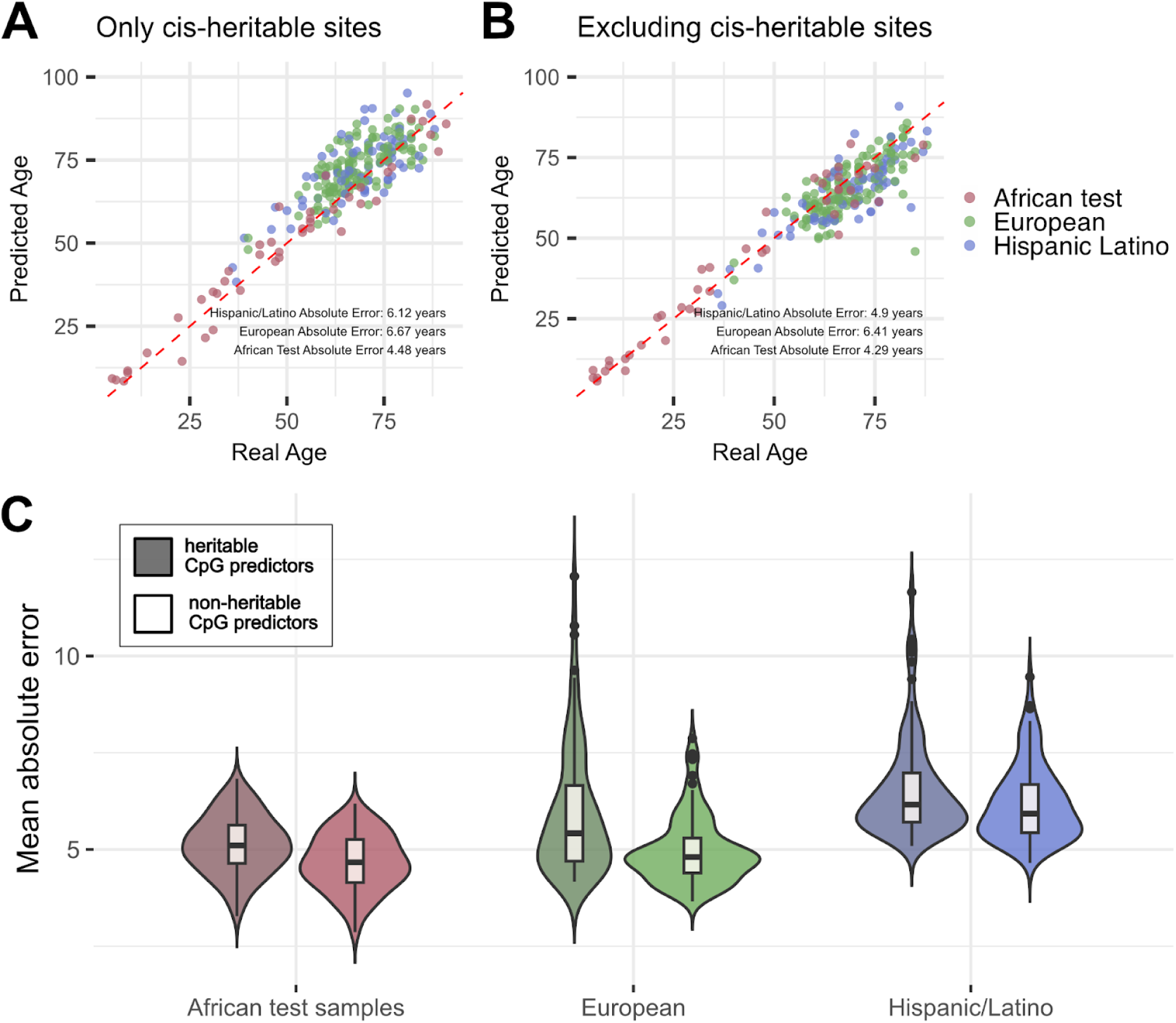
Performance of epigenetic clocks trained on a set of CpG sites that do not exhibit significant *cis*-heritability of DNA methylation and a set of CpG sites that do exhibit evidence of *cis*-heritability. The top row of panels show predicted age on the y-axis plotted against true age on the x-axis for the epigenetic age prediction models that are comprised of either significantly *cis*-heritable CpG predictors (A) or not significantly *cis*-heritable predictors. Panels A and B are each based on a single model that exhibited median accuracy among a total of 100 models. Panel C depicts the mean absolute error across all 100 models within the African test subset, European, and Hispanic/Latino cohort, restricted to individuals aged 36-91. Models based on CpG predictors that are not significantly impacted by *cis*-genetic variation exhibit lower absolute error and less bias when applied out-of-cohort than models based on CpG sites that are significantly heritable.

### The combined effects of age-associated meQTL correlate with age and epigenetic age acceleration

Looking beyond age prediction, we wondered if meQTL variation at age-associated CpG sites had biologically meaningful consequences for aging and longevity. To this end, we developed genotype-based epigenetic aging scores (EAS), which sum up the effects of meQTL variants on DNA methylation, weighted by the effect of DNA methylation on age; the result is analogous to a polygenic score that captures an individual genome’s total burden of epigenetic age-elevating variants (Supplementary Figure 8). For each cohort, we build an EAS model based on CpG sites that are both associated with age in that population’s EWAS at a relaxed significance threshold (p < 0.001) and are significantly influenced by a *cis*-genetic variant in that population’s baseline meQTL scan (see Methods). Our ‡Khomani San model was based on 668 SNPs near 718 distinct CpGs, the Baka model on 987 SNPs near 1075 CpGs, and the Himba model on 1921 SNPs near 1995 CpGs. We then applied each of these models to genotype data from individuals within that cohort. Interestingly, we found that older individuals tended to have lower EAS, and consistently observed an overall negative relationship with age across all comparisons (Figure 7A). Based on this observation, we hypothesize that having a lower burden epigenetic age-elevating genetic variants might enable individuals in these populations to achieve greater longevity.

**Figure 7.**
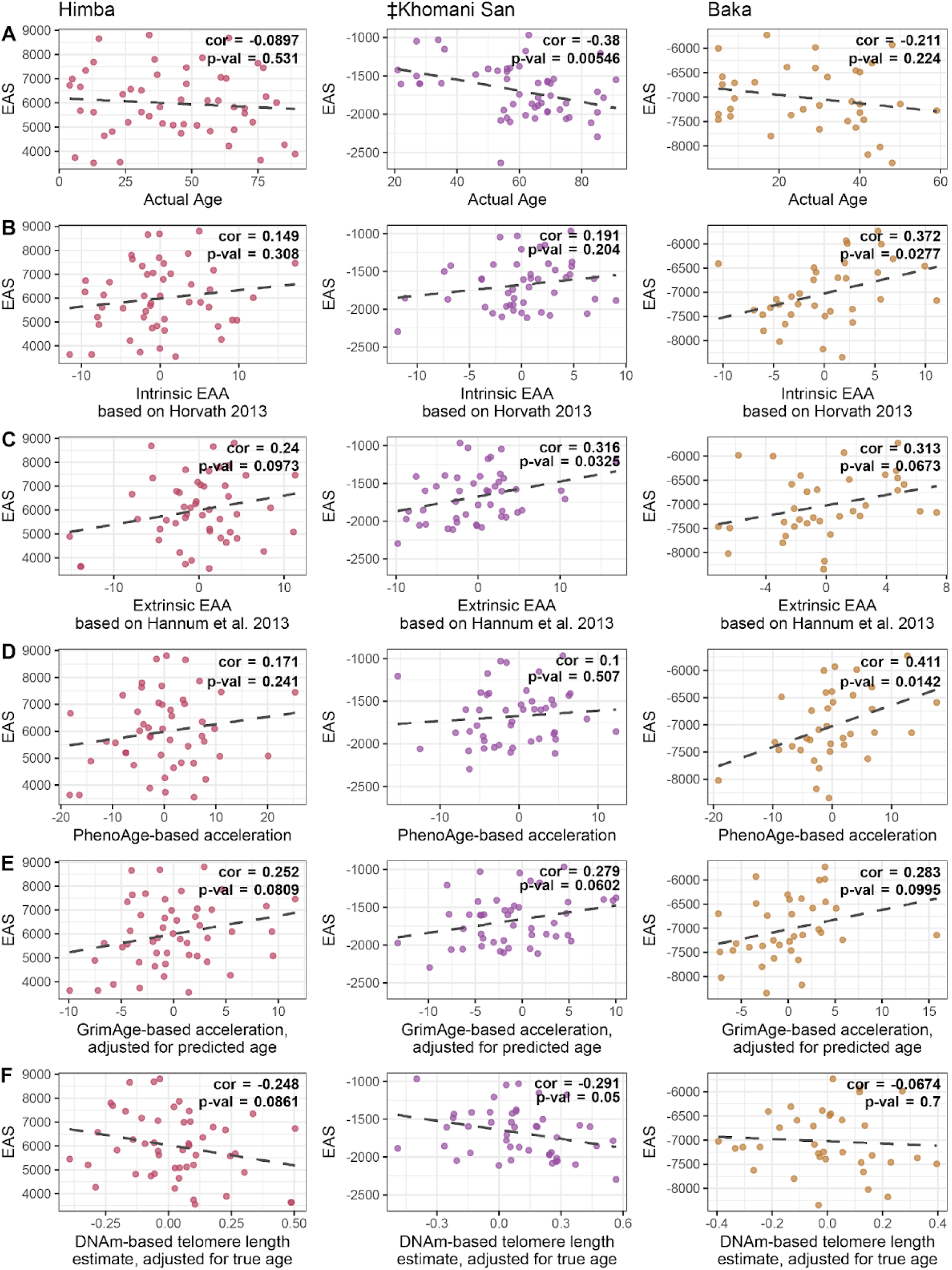
Relationship between EAS and aging metrics. Scatterplots show the relationship between the genotype-based epigenetic aging score (EAS) and various metrics of age for the Himba, ≠Khomani San, and Baka. Each EAS model was built from the respective population’s epigenome-wide association study and baseline meQTL scan results. Individuals’ EAS values were plotted against A) chronological age itself, B) ‘Intrinsic Epigenetic Age Acceleration’, based on the Horvath multi-tissue age predictor, C) ‘Extrinsic Epigenetic Age Acceleration’, based on the Horvath multi-tissue age predictor, D) epigenetic age acceleration based on PhenoAge, E) epigenetic age acceleration based on GrimAge, adjusted for predicted age, and F) DNA methylation-based telomere length, adjusted for true age.

Seeking additional evidence to address this hypothesis, we compared these genotype-based EAS values with the several published measures of epigenetic age acceleration that are based solely on DNA methylation data^20,21,65,71^. These epigenetic age acceleration metrics have been shown to be associated with various aging related phenomena^16,71–73^.

Interestingly, we found associations between EAS and multiple measures of biological aging or accelerated epigenetic aging; EAS trends towards being positively correlated with measures of ‘intrinsic’ and ‘extrinsic’ epigenetic age acceleration^71^, while it trends towards being negatively correlated with an age-adjusted DNA methylation-based estimate of telomere length^65^ (Figure 7B-F, Supplementary Figure 8). While not always significant, the trends we observe are consistently in the expected direction across populations and across acceleration metrics, supporting the role of genetic variants in influencing the pace of biological and epigenetic aging^13,14,52,74^.

## Discussion

As a result of a growing interest in using epigenetic age predictors in clinical settings^75–77^, models such as FitAge, PhenoAge, and GrimAge have been explicitly trained to capture traits such as maximal oxygen uptake (VO_2_max), healthspan, and lifespan, respectively^20–23^. Epigenetic clocks intended for forensic applications, on the other hand, are concerned with accurately predicting an individual’s chronological age, regardless of lifestyle or overall health. However, whatever the goals of an epigenetic clock, relatively little attention has been paid to the issue of transferability; i.e. how well a predictive model trained in one population or cohort performs when applied in a genetically diverged population^36,78^. Researchers have been grappling with an analogous issue in the development of polygenic risk scores (PRS), models that predict an individuals’ risk of complex disease based on their genotype. While initially heralded as a promising tool that would enable personalized genomic medicine, recent work demonstrates that applying PRS out-of-cohort can actually worsen health disparities due to poor transferability across human populations^40,79–81^. Some of the underlying mechanisms that account for PRS’ lack of generalizability, such as differences in allele frequencies and trait heritability across populations may also be relevant for epigenetic clocks. In addition to these concerns, however, a loss of epigenetic clock transferability could also be driven by variation in the lifestyle or environmental factors across populations, which can directly impact DNA methylation patterns without perturbing the underlying DNA sequence.

In testing several popular epigenetic clocks on three diverse African cohorts, we find that almost all exhibit significant among-population differences in prediction error, even after accounting for differences in data missingness, cohort age ranges, and potential tissue-predictor mismatch (see Methods). Only the Horvath 2013 predictor showed no significant differences in age-adjusted error among cohorts (Figure 1A). This differs from previous results from our group and others that found that the Horvath 2013 predictor produces systematically different estimates for African-ancestry and Hispanic/Latino individuals compared to European-ancestry individuals^35,66,82^. This discrepancy may be due in part to differences in implementation; in our prior work, we applied the Horvath algorithm to quality control filtered DNA methylation values that had been pre-normalized and imputed^66^, whereas here it was applied to raw and unfiltered data according to the online platform standards (see dnamage.clockfoundation.org).

Unlike previous work, we also did not find any significant differences among populations for any of the epigenetic age acceleration metrics derived from the Horvath and Hannum clocks^20,32–35^ (Supplementary Figure 5). It has been suggested that apparent among-population variations in epigenetic age acceleration could indicate, or help explain, real differences in the average health and/or longevity of different human groups^20,25,32–35^. However, contradictory results across clocks and across studies, along with decreasing prediction accuracy in genetically diverged samples, suggest underlying issues in clock transferability^20,32–34,37–39,83^. Since most epigenetic clocks have been trained primarily on European-ancestry individuals living in industrialized societies, we suggest that these discrepancies might be partially explained by differential transferability across cohorts.

Although we did not have the health and mortality data required to rigorously evaluate the relationship between epigenetic age acceleration and health outcomes in the Himba, ‡Khomani San, and Baka, our results demonstrate that poor transferability across ancestries should be considered more critically as a possible explanation for among-population differences. For example, we found that PhenoAge and GrimAge2 exhibited qualitatively different patterns in our among-population comparisons, even though both of these epigenetic clocks were designed to capture signatures of age-associated morbidity and mortality; relative to European-ancestry individuals, PhenoAge significantly underestimates individual Himba age, while GrimAge2 using true age significantly overestimates Himba age, and GrimAge2 using predicted age finds no significant difference (Figure 1E,G,H). Although these two clocks were designed to capture slightly different aspects of the aging phenotype, we would not expect these kinds of inconsistencies if Himba individuals are truly aging faster (or slower) on an epigenetic level compared to European-ancestry individuals.

Furthermore, we find that ‘cryptic’ meQTL variation is a significant factor affecting age association and epigenetic clock performance. We rescue hundreds of age associations after accounting for meQTL genotype and find that all of the published epigenetic clocks we tested included a large proportion of CpG predictors that are impacted by *cis* genetic variation, of which a substantial fraction is rare or absent in European-ancestry populations. Populations that experienced the out-of-African migration bottleneck carry only a subset of the genetic variation that exists in African populations^84^; therefore, we expect that our analyses capture most of the meQTL variation that exists in European and Hispanic/Latino ancestry populations. In training our own epigenetic clocks, we show that excluding CpG sites with detectable *cis*-heritability improves prediction accuracy and reduces bias when applied out-of-cohort relative to clocks that only include heritable CpG predictors (Table 1, Figure 6). Until we have a better understanding of the genetic architecture of DNA methylation variation across diverse human populations, this seems the most feasible approach to improving transferability. This is particularly true for forensic applications, where generating an accurate estimate of chronological age regardless of health status is the primary goal. However, given the remarkably strong correlation of meQTL effect sizes across genetically diverged populations (Figure 3A-C), future epigenetic clocks could see even greater improvements by explicitly accounting for individual genetic variation. Therefore, while environmental differences could still drive variation in DNA methylation, and thus prediction error, across cohorts, we show that heritable factors a significant role in transferability.

Although we refer to the difference between individual chronological age and predicted age based on some epigenetic model as ‘prediction error’ throughout this paper, it must be noted that these deviations are not necessarily true errors; for some epigenetic clocks, these deviations do appear to reflect meaningful variation in human health, morbidity, and/or mortality within specific populations^75–77^. However, as we have outlined above, it is not clear to what degree among-population differences in mean estimates are indicative of genuine variation in the rate at which different human populations age versus a simple lack of transferability. Answering this question will require a complete understanding of the connections between various genetic, environmental, and lifestyle factors and CpG methylation, as well as their interactions and downstream effects on the aging phenotype. Our work here focuses on the genetic factors, whose effects on CpG methylation we are able to dissect by jointly analyzing DNA methylation and genotype data. Additionally, our multi-population study design enables us to characterize the extent to which the genetic architecture of age-associated CpG methylation is shared across diverse genetic backgrounds and environments (Figure 3).

We were also able to investigate the potential impact of genetic variation on the aging phenotype by developing an epigenetic aging score (EAS) that reflects the cumulative effect of meQTL variants that influence age-associated CpGs. We find suggestive evidence that our EAS is associated with older chronological age and with epigenetic age acceleration in these populations (Figure 7). If genetic factors influence lifespan and healthspan, we might expect that older individuals will have lower EAS (i.e. a lower burden of epigenetic age increasing variants) while younger individuals will exhibit a wider range of EAS values. In our data, we find that this pattern manifests as a slight negative correlation between EAS and chronological age. Furthermore, we find that EAS is also correlated with various estimates of epigenetic age acceleration, despite the fact that these metrics were not trained on African populations and thus are likely underpowered. Although not always significant, the consistency of these associations in the expected direction is nevertheless compelling. These results also corroborate previous work that has found that both healthspan and lifespan are heritable, polygenic traits^13,14,52,74,85^. Horvath 2013 noted 21 genes that carried common variants associated with increased epigenetic age in his multi-tissue clock. Interestingly, six of these genes (FAM123C, LEPR, CHD7, CTNND2, TMEM132D, and MACF1) were represented in at least one of our population-specific EAS models. These results suggest that our EAS approach is picking up on real signals of a genetic predisposition to accelerated biological aging within these African populations that warrant further investigation. Although our sample sizes are small, there is reason to think that our study is particularly well-suited to identify these signals, especially in the association between EAS and age. Among-individual variation in socio-economic status, diet, and other lifestyle factors is relatively low within the Himba, ‡Khomani San, and Baka compared to most industrialized populations, which would allow the influence of genetic variation to be more readily detectable.

The extent to which the pace of epigenetic aging is determined and modulated by heritable versus non-heritable factors is still very much an open question, with important implications for the problem of transferability. These issues are becoming more relevant as epigenetic clocks are being more frequently applied in contexts where these genetic and environmental factors are often confounded. For example, the relatively new subfield of ‘social epigenomics’ seeks to understand how socio-economic and environmental factors influence DNA methylation and drive health disparities in cosmopolitan populations^25^. Differences in epigenetic age acceleration among racial and/or ethnic groups are typically interpreted as arising from systemic differences in socioeconomic status, etc. However, it is possible that poor model transferability partially accounts for these observations. This alternative explanation does not minimize the growing body of evidence that has broadly demonstrated that various social determinants of health, such as psychosocial stress^86^, diet^87^, and smoking behavior^88^ influence DNA methylation. Rather, we caution that genetic ancestry should be more carefully considered in studies of epigenetic aging and its consequences for human health, as it is often confounded with underserved minority status, particularly in the Global North.

## Methods

### DNA methylation microarray quality control and filtering

DNA was bisulfite converted, whole-genome amplified, fragmented, and hybridized to the Illumina Infinium HumanMethylation450 (> 485,000 CpG sites) BeadChip array for the Baka and ‡Khomani San samples and the EPIC Array (> 845,000 CpG sites) for the Himba samples. DNA methylation array data was generated in 4 batches with both the ‡Khomani San and Himba samples separated across two batches (Supplementary Table 6). One ‡Khomani San individual was typed twice across both batches and two Baka individuals were typed twice in the same batch. The overall intra-class correlations between DNA methylation values for these 3 sets of replicates were 0.9985, 0.9991, and 0.9989, respectively. One Himba individual, sampled three years apart, was typed across the two Himba batches. The overall intra-class correlation between methylation values from this individual was 0.9974, lower than for a purely technical replicate, as expected. Only the first sample from this individual was used in the EWAS and meQTL analyses. One Baka individual was flagged for having abnormally low bisulfite controls and removed from further analyses.

We removed DNA methylation probes with a detection p-value > 0.01 in greater than 5% of samples, as well as any probes that have been reported to be cross-reactive, map to multiple regions, or to the sex chromosomes^89,90^ (Supplementary table 7). Any remaining values with detection p-values > 0.01 were set to NA. We also removed CpG sites that were likely to be impacted by SNPs in or near the probe sequence in a population specific manner using our previously published software, probeSNPffer^91^. Specifically, we retrieved the hg19 genomic coordinate of the target cytosine for each DNA methylation array probe and searched the full 50 base pair probe region, the next base extension (for type 1 probes), and the extension base (for type 2 probes) for overlap with SNPs segregating at > 5% frequency in a given population^91^.

SNPs within array probes can lead to reduction in probe hybridization efficiency and unreliable methylation signal^89,92,93^. An additional 27,242, 9,254, and 61,662 probes were pruned from the Baka, merged ‡Khomani San, and merged Himba DNA methylation datasets, respectively from this step.

After these filtering steps, we were left with 713,988 CpG sites in the Himba dataset, 418,629 sites in the ‡Khomani San dataset, and 400,893 sites in the Baka dataset. There was an overlap of 355,103 sites across all three populations that we used for combined analyses. DNA methylation values were background and color corrected, and technical differences between type 1 and type 2 probes were corrected by performing BMIQ normalization using the wateRmelon^94^ and minfi^95^ R packages. All analyses were performed using continuous DNA methylation beta values for each CpG site, which range from 0 (indicating that the site is completely unmethylated) to 1 (completely methylated).

### Genotype data quality control and filtering

Genotype data was generated using multiple arrays for ‡Khomani San and Himba samples, while the 35 Baka individuals were all genotyped on the Illumina OmniOne array (Supplementary table 8). The Baka dataset was known to contain 9 family trios and 9 unrelated individuals. All genotype data were oriented to match the 1000 genomes Phase 3 GRCh37 reference, filtered to exclude SNPs with a genotype missing rate > 5%, minor allele frequency of < 1%, and Hardy-Weinberg deviation p-value <0.0001. We removed all indels and A/T or C/G transversion variants. Sample sizes and pre-imputation variant counts are listed in Supplementary Table 8.

Each genotype array dataset was phased using SHAPEITv2.r790^96^ and imputed using the Positional Burrows-Wheeler Transformation (PWB)^97^ to the African Genomics Resources Panel (89,838,088 autosomal variants, 4956 samples) via the Sanger Imputation Service^53^. We assessed imputation accuracy in our samples by calculating imputed genotype concordance. For the ‡Khomani San, we compared imputed genotypes for 37 individuals with whole exome sequence data. For the Himba, we compared genotypes imputed from MEGAex array data with genotype calls uniquely typed on the H3Africa array data for 3 Himba individuals genotyped on both platforms. The overall concordance with the ‡Khomani San exome variants was 95.7-97.7% across all genotype arrays for variants of any impute quality INFO score (Supplementary Figure 9D). The average concordance of imputed H3Africa SNPs was 98% for the 3 Himba individuals typed on both H3Africa and MEGAex (Supplementary Figure 10D). We also stratified concordance by imputed quality INFO score, and observed 99% concordance across all genotype arrays in the ‡Khomani San and Himba at a >0.95 INFO score (Supplementary Figure 9D, 10D). This observation informed our choice to only retain imputed variants with >0.95 INFO score for subsequent analyses. Imputed data from OmniExpress and then MEGAex arrays performed slightly better on concordance metrics for all INFO score bins than 550K array (Supplementary Figure 9A-D), most likely due to denser genotyping, so OmniExpress and MEGA array genotype data were used for the 17 ‡Khomani San individuals typed on multiple arrays. After this filtering and merging across genotype arrays we retained 66,484,843 high-quality autosomal variants for the ‡Khomani San, 78,738,543 for the Himba, and 75,739,815 for the Baka.

### Epigenome-wide association studies (EWAS)

We used EMMAX^98^ with the dosage option to test for the association between age and methylation level separately in each population, accounting for population-specific scaled covariates and a Balding-Nichols kinship matrix (Eq 1-3).

DNA methylation array data are known to exhibit significant batch effects; that is, samples on one run vary systematically from samples on another due to technical artifacts. We controlled for DNA methylation array batch effects by including the first 20 PCs of control probe intensities^99^ as covariates in the EWAS for the Himba (Eq 1). Regressing out these control probe PCs eliminates batch effects in the first two methylation PCs (Supplementary Figure 11C-D). We did not have access to the raw intensities for the Baka and ‡Khomani San methylation datasets so we controlled for technical artifacts by including batch number as a covariate in ‡Khomani San where samples were split across batches (Supplementary Figure 11A-B). To control for technical artifacts present within a batch we included the combination of the first 5 DNA methylation PCs that we found best reduced genomic inflation. We did not include all of the first 5 methylation PCs as covariates to mitigate power loss as up to one third of the methylome has been found to show association with age^2^.

We included sex and the first 5 genetic PCs as covariates in all models (Eq 1-3). We computed the first 5 genetic PCs on LD pruned (PLINK1.9^68^ --indep-pairwise 50 5 0.3) variants above 5% frequency within each population to include as covariates. There was no evidence of clustering based on genotype data in the ‡Khomani San (Supplementary Figure 12). We estimated cell type proportions using the R package EpiDish^63^, leveraging DNA methylation data from a reference panel of 12 different blood cell types, epithelial cells, and fibroblast cells. The proportions estimated by this method correspond closely to previous estimates of saliva cell composition^64^ (Supplementary Figure 13). As neutrophils and epithelial cells together account for nearly 100% of the cells in our saliva samples, we included just the neutrophil proportion as an additional covariate in our models. Cell type proportion estimates for the replicate samples were highly similar (Supplementary table 9). We also performed reference-free estimates of cell type proportions using the TOAST^100^ R package. Under a k=2 cluster model, the correlation with the reference-based neutrophil and epithelial cell proportion estimates was 0.98.

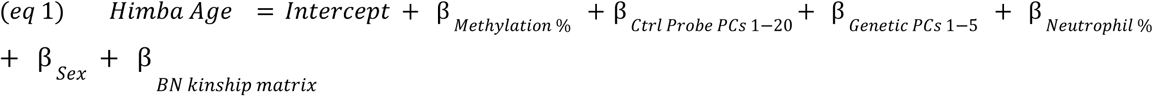

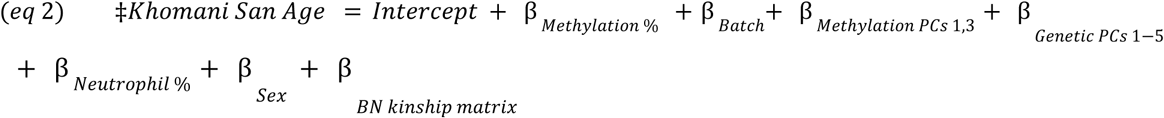

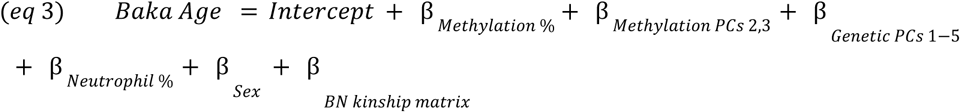

We used the metagen^101^ R software package to conduct a fixed-effect meta-analysis of our EWAS results from all three populations. Significance was determined at a Bonferroni corrected *p*-value of 0.05, correcting for the number of overlapping CpGs across the three populations. We set the Hartung and Knapp adjustment to false and the between-study variance method to REML.

### Baseline *cis-*meQTL scan

We used EMMAX^98^ with DNA methylation value as the dependent variable to identify *cis*-variants that are significantly associated with DNA methylation levels at each CpG site. SNPs with a minor allele count of less than 2 were removed to leave 2,432,803 for the ‡Khomani San, 6,594,680 for the Himba, and 4,944,508 for the Baka. We performed within-population *cis*-meQTL scans using LD-pruned genotype datasets (generated using the PLINK1.9^68^ option --indep-pairwise 50 5 0.5) by testing each SNP within a 200kb window (100kb upstream and downstream) of the target CpG for association with DNA methylation level. The same population-specific scaled covariates as in the EWAS scan were used with the addition of age (eqs 4-6). We determined significance at a p-value of 0.05 corrected for the number of SNPs tested at each CpG.

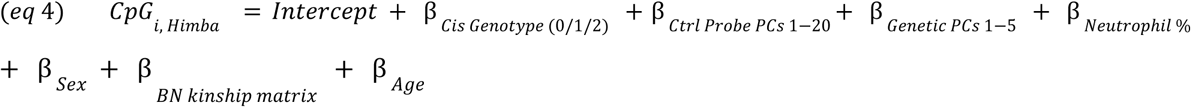

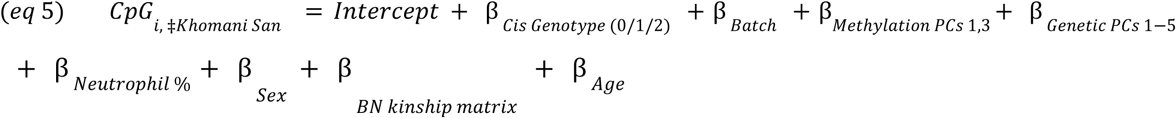

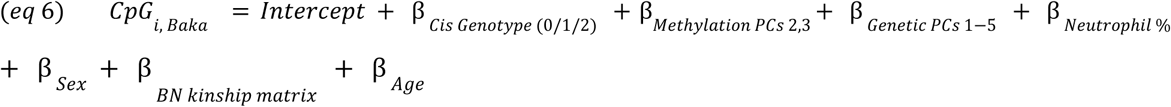

### Heritability of CpG methylation

We estimated *cis*-heritability of DNA methylation at each CpG site using GCTA^102^ within FUSION^67^ and default parameters (--reml --reml-no-constrain --reml-lrt 1). We tested a 1Mb window (i.e. 500kb upstream and downstream) around each CpG site. We used the same genetic datasets as used in the baseline meQTL scan prior to the LD pruning step. The same covariates were used as in the baseline meQTL scans (Eq. 4-6).

### FUSION *cis*-meQTL scan

We modified functions from the FUSION^67^ software package, originally designed to uncover the *cis* genetic architecture of gene expression, to test elastic net, LASSO^68^, SuSie (sum of single-effect)^69^, and best single meQTL regression models to explain methylation levels at each CpG site. The former 3 regression models allow for multiple SNP effects to jointly influence methylation rather than testing the effect of each SNP independently as in our baseline scan.

Only CpGs with significant (p-value < 0.05) *cis*-heritabilty were modeled using the 4 regression models. The FUSION framework conducts 5-fold cross-validation analyses to select the regression model that yields the highest R-squared in explaining *cis* genetic variation’s effect on DNA methylation and stores the effect sizes (i.e., weights) associated with each variant under each model.

### MeQTL-adjusted EWAS

We re-ran our EWAS (Eq 1-4), this time testing for age associations with the residual values after regressing out the top meQTL genotypes from respective CpG’s methylation values. This was done for each CpG site with a significant meQTL association. We used the same covariates as the original EWAS (Eq 1-3).

### Testing published epigenetic clocks

We tested all 10 published age predictors available through the Clock Foundation online portal at dnamage.clockfoundation.org. The Horvath^13^, Hannum^14^, Skin and blood^54^, and Zhang elastic net^55^ clocks are all chronological age predictors built using penalized linear regression. The PhenoAge^20^, GrimAge^21^, GrimeAge2^22^, and Fitage^23^ clocks are built on CpGs associated with surrogate measures of biological age, capturing variables that predict lifespan, healthspan, and mortality risk. GrimAge and GrimAge2 models incorporate chronological age within their surrogate measure and can be constructed using actual chronological age (GrimAge on true age), or using estimates of chronological age from the Skin and Blood clock predictor^54^(GrimAge on predicted age). See eTable 1 in Krieger et al., 2024^31^ for detailed descriptions of each predictor.

Each model’s age predictions are a weighted sum of an individual’s DNA methylation values at the predictor CpG sites, and are thus very sensitive to missing data. Therefore, in order to fairly compare predictions across populations, and in accordance with recommendations published with the online tool, we uploaded raw, unfiltered beta values. Additionally, we restricted our dataset to CpG sites that are common to both EPIC and 450k DNA methylation arrays. The online tool also performs a host of additional analyses based on the input DNA methylation values, including an unpublished tissue type prediction algorithm. We only evaluated performance for samples predicted to be saliva, blood PBMC, or whole blood, as we found that the estimated cell type proportions for these three tissue types were essentially indistinguishable in our dataset (Supplementary Figure 14). Our final sample sizes for these clock validation analyses were: ‡Khomani San (n = 46), Himba (n = 49), Baka (n = 35), Hispanic/Latino (n = 69), European (n = 130). We adjusted prediction errors by regressing out chronological age before evaluating among-population differences.This was necessary to avoid confounding based on the different age distributions within our cohorts because prediction accuracy can vary systematically by age for some epigenetic clocks^56^. By taking these steps, we ensured that the differences in prediction error are not due to sampling design, tissue type or other technical issues. We identified significant among-population differences in the distribution of age-adjusted prediction errors by ANOVA, followed by a Tukey test to identify significant pairwise differences.

We also assessed the differences in epigenetic age acceleration (EAA) metrics across cohorts (Supplementary Figure 3). The Horvath residual, Hannum residual, GrimAge and PhenoAge acceleration metrics are calculated by adjusting the epigenetic age estimated by each predictor for chronological age. Intrinsic epigenetic aging acceleration (EAA) measures the component of EAA that is not influenced by changes in white blood cell count with age (i.e., it is the residual of the Horvath estimate after regressing out both chronological age and DNA methylation-based estimates of blood cell proportions). Extrinsic EAA instead captures both this intrinsic component and age-related changes in white blood cell composition by using the residuals of an enhanced version of the Hannum-based age estimate after regressing out chronological age^71^. Positive values of these measures indicate that an individual’s predicted age is higher than their actual chronological age. The DNA methylation-based telomere length acceleration estimate is generated by regressing out chronological age from DNA methylation-based estimates of telomere length^65^. Positive values indicate that an individual is estimated to have longer telomeres than would be expected for their age^103^. Telomeres tend to get shorter with age, and in association with increased risk of age-related diseases^104,105^.

### Constructing chronological age predictor

We developed chronological age prediction models using elastic net regression. We selected predictors using the cv.glmnet function in R, employing leave-one-out cross-validation on the training dataset. We conducted 100 different splits of our dataset into training and test sets to construct the heritable and non-heritable prediction models. Each training dataset was created from randomly sampling 70% of the ‡Khomani San and Himba samples along with 63% of the Baka samples. For the trios contained in the Baka dataset, children were never included with their parent(s) for training. Our heritable models were built from 123,194 possible predictor CpGs found to be significantly heritable (p < 0.05) or influenced by meQTL in our baseline scan in any of the three populations. Our non-heritable models were built from 227,770 possible predictor CpGs not found to be significantly heritable and not influenced by meQTL in our baseline scan in any of the three populations. We conducted a grid search to optimize the alpha parameter for the elastic net regression model. Alpha values ranged from 0 to 1 in increments of 0.05. For each alpha, we used leave-one-out cross-validation on the training data to construct the model and selected the lambda value that minimized the mean squared error (MSE). The best-performing model was then identified based on the alpha value that resulted in the lowest MSE on the held out test dataset. We used transformed chronological ages following Horvath’s method ^13^ to account for the logarithmic relationship observed at many sites between methylation and age in children and young adults. We then applied each of our 100 heritable and 100 non-heritable epigenetic age prediction models to the out-of-sample

European-ancestry and Hispanic/Latino datasets after first normalizing these data using the wateRmelon^94^ package’s BMIQ function to match our normalization procedures for the African DNA methylation data.

### Epigenetic aging score (EAS) models

We constructed an EAS for each African population by using the baseline meQTL results to identify SNPs that have a strong influence on DNA methylation levels at a nearby CpG site, retaining only the most significant SNP per CpG site. We intersected this list with each populations’ meQTL-regressed EWAS results to identify CpG sites that are both influenced by *cis*-meQTL and age-associated, using a relaxed significance threshold of 10^-3^ for the latter. We extracted the effect of each SNP allele on CpG methylation and CpG methylation on age to construct the EAS, which is effectively a polygenic risk score that represents the total burden of epigenetic age increasing genetic variants on an individual’s genome *i* (Eq. 7, Supplementary Figure 14). The effect of each SNP *j* on age is given by its effect on DNA methylation at the corresponding CpG site, weighted by the effect of DNA methylation level at that CpG site on age. This weighting ensures that the direction of the SNP-on-age effect is consistent across loci, which are then summed to yield an EAS value.

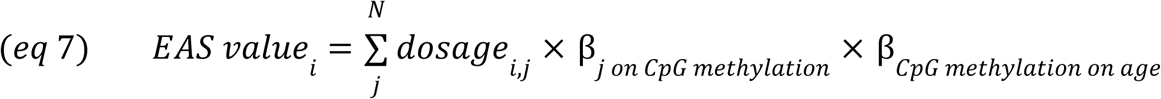

## Acknowledgements

We would like to thank the Working Group of Indigenous Minorities in Southern Africa and the South African San Institute for their advice regarding community research. We thank the ≠Khomani San, Baka, and Himba communities in which we have sampled; without their support, this study would not have been possible. B.M.H and G.L.M. were supported by the National Institutes of Health grant R35GM133531. S.G. was supported by the National Institutes of Health Center of Biomedical Research Excellence grant 1P20GM139769 and the National Institute of Justice grant 2016-DN-BX-0011. B.S. was supported by the National Science Foundation grant BCS-1534682. The content is solely the responsibility of the authors and does not necessarily represent the official views of the National Institutes of Health, National Science Foundation, or National Institute of Justice. The funders had no role in the decision to publish or prepare the manuscript.

## Author Contributions

S.G. and G.L.M. conducted analyses. S.G. and G.L.M. wrote the manuscript with input from all authors. B.S., S.P., E.P., A.F., M.F., L.Q. contributed data for analyses. H.A. conducted preliminary analyses on newly generated data. S.G. and B.M.H. conceived and supervised the study.

## Competing Interests

The authors have no competing interests to declare.

## Materials and Correspondence

Correspondence should be directed to Shyamalika Gopalan

## Data Availability

The data from the Baka and ≠Khomani San used in this article have been previously submitted to the European Genome-Phenome Archive (EGA) (www.ebi.ac.uk/ega/home) and GEO (https://www.ncbi.nlm.nih.gov/geo). The SNP and methylation array data for the Baka can be found under the EGA accession numbers EGAS00001001066 and EGAS00001002226. The SNP and methylation array data for the ≠Khomani San can be found under the GEO super series GSE99091. SNP array data for the Himba are available via dbGaP, accession <u>phs001995.v1.p1</u>. The newly generated Himba methylation data will be available via GEO deposition. The 1000 Genomes Phase 3^1^ data can be accessed via http://ftp.1000genomes.ebi.ac.uk/vol1/ftp/phase3/.

## Notes

### Competing Interest Statement

The authors have declared no competing interest.

