## Supplement for "Common DNA sequence variation influences epigenetic aging in African populations"

**Table of Contents**

**Supplementary Figures	2**

Supplementary Figure 1 - Flow chart of study design	2

Supplementary Figure 2 - Predicted age versus chronological age from published epigenetic clocks	3

Supplementary Figure 3 - Correlation of DARC genotype status with estimated neutrophil proportion	4

Supplementary Figure 4 - Correlation of DARC genotype status with prediction error across epigenetic clocks	5

Supplementary Figure 5 - Distributions of epigenetic age acceleration estimates across diverse cohorts	6

Supplementary Figure 6 - Overlap of significant CpG sites across the Himba, ‡Khomani San, Baka	7

Supplementary Figure 7 - Genomic Inflation comparisons for meQTL-adjusted vs unadjusted EWAS	8

Supplementary Figure 8 - Relationship between EAS and metrics of epigenetic age acceleration	9

Supplementary Figure 9 - Concordance of imputed genotypes with exome calls by INFO score.	11

Supplementary Figure 10 - Concordance of imputed genotypes with independent calls from genotyping on the MEGAex array by INFO score	12

Supplementary Figure 11 - Principal component analysis of batch effects in DNA methylation data	13

Supplementary Figure 12 - Principal components analysis of genotype data	14

Supplementary Figure 13 - Estimated cell type composition by tissue type and population	15

Supplementary Figure 14 - Schematic of the epigenetic aging score	16

**Supplementary Tables	17**

Supplementary Table 1	17

Supplementary Table 2	18

Supplementary Table 3	19

Supplementary Table 4	20

Supplementary Table 5	21

Supplementary Table 6	22

Supplementary Table 7	23

Supplementary Table 8	24

Supplementary Table 9	25

### Supplementary Figures

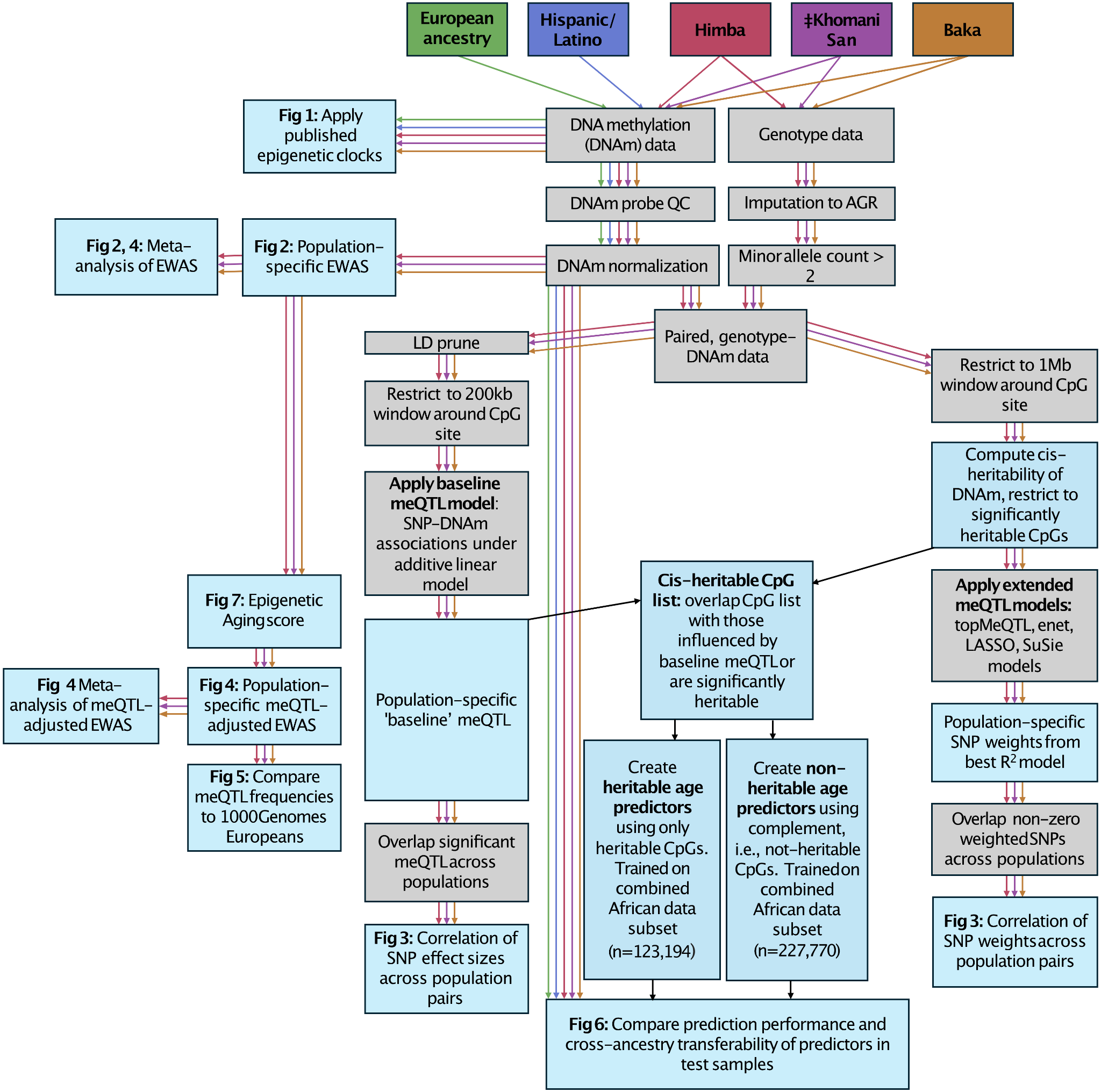

#### Supplementary Figure 1 - Flow chart of study design

Grey boxes indicate analysis steps and blue boxes indicate results. Colored arrows indicate steps that are done in parallel, separately in multiple populations, while black arrows indicate steps that are done on combined data or combined results.

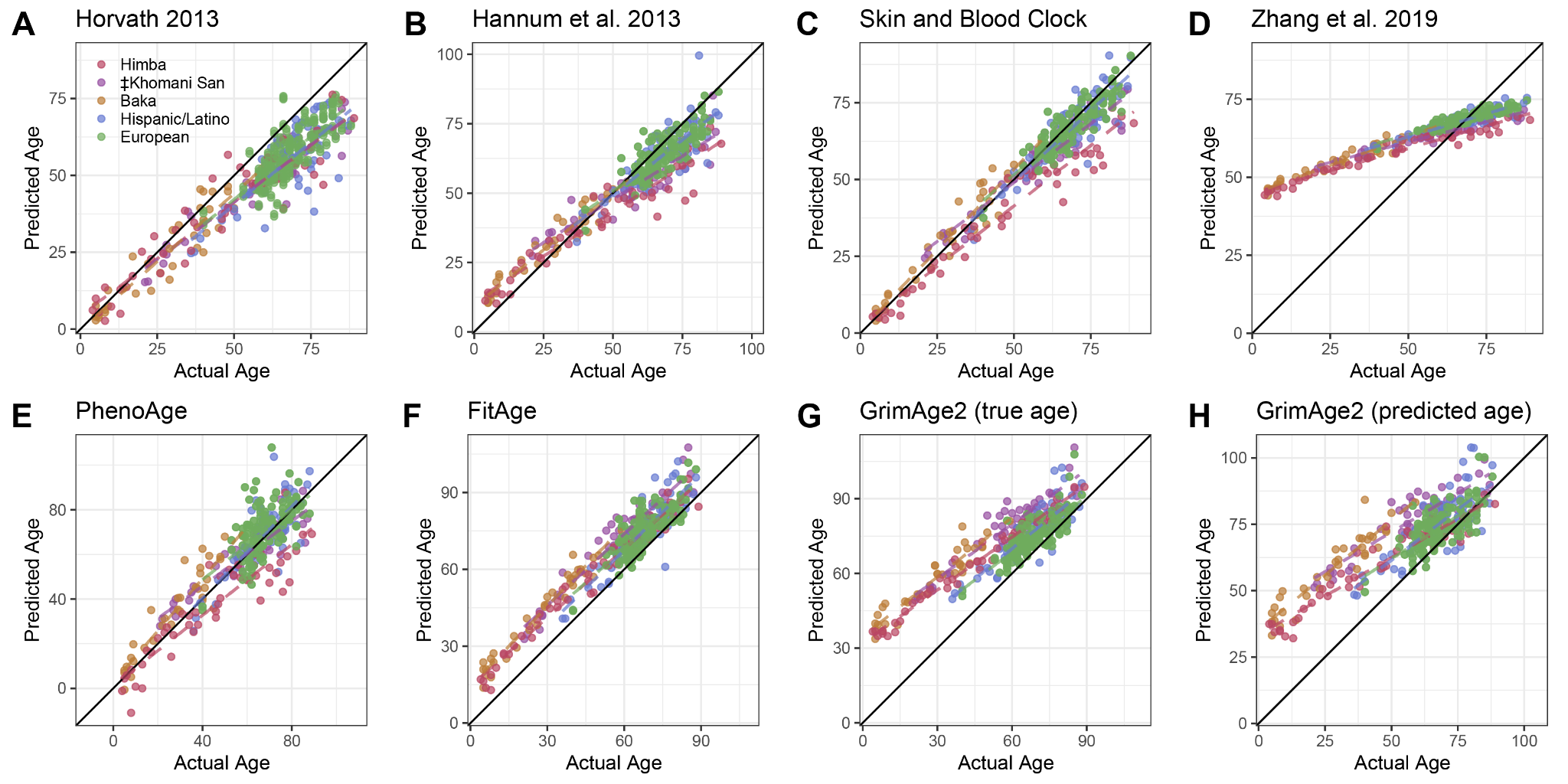

#### Supplementary Figure 2 - Predicted age versus chronological age from published epigenetic clocks

Each point represents one individual for whom DNA methylation data was available. Reported chronological age is plotted on the x-axis, while the age predicted by the epigenetic clock is plotted on the y-axis. Results from the first version of GrimAge are not shown here for simplicity, as both versions produced very similar results.

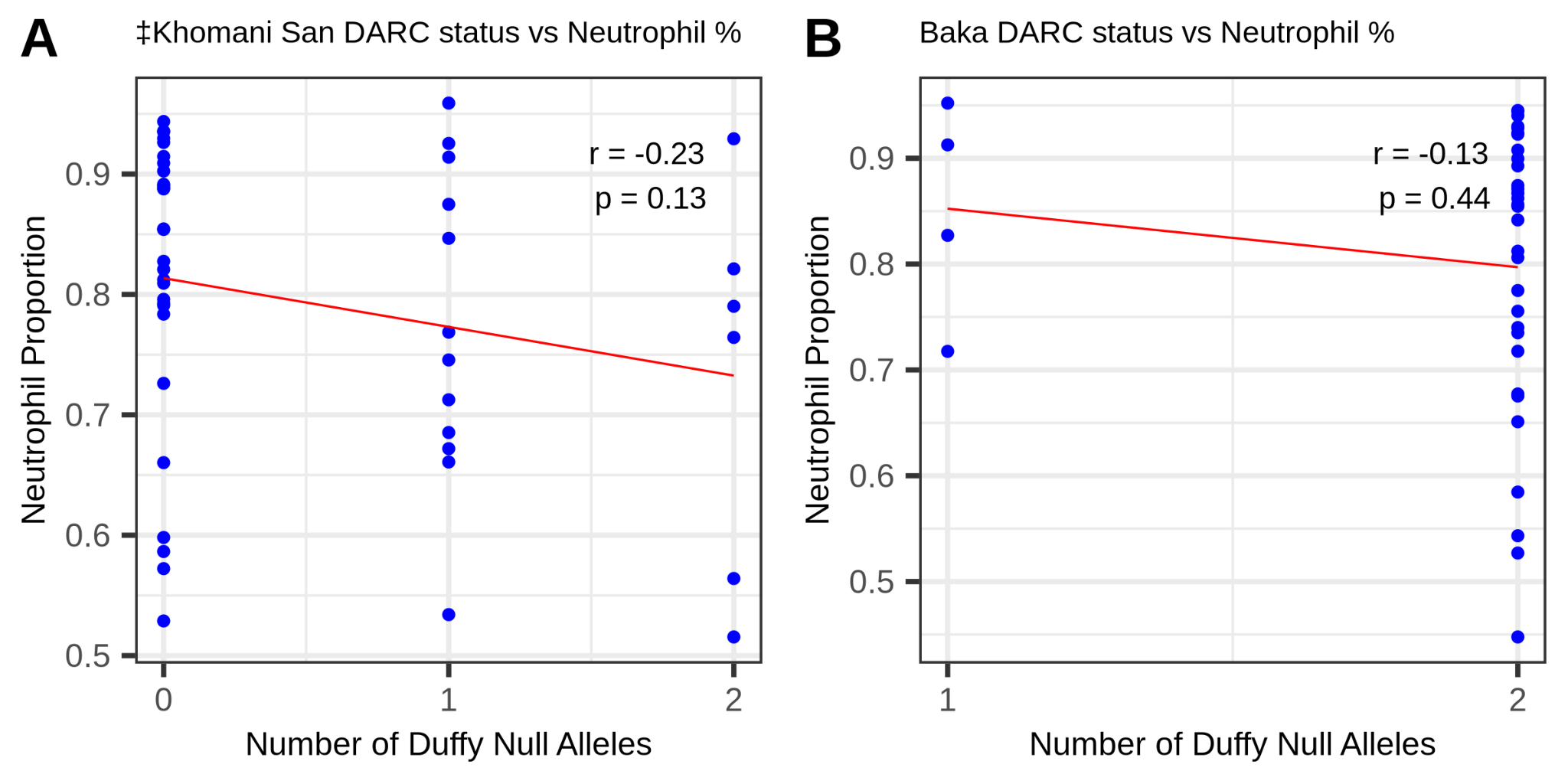

#### Supplementary Figure 3 - Correlation of DARC genotype status with estimated neutrophil proportion

The number of Duffy null alleles carried by an individual is plotted on the x-axis against their proportion of neutrophils, as estimated by EpiDish, on the y-axis for the ‡Khomani San (A) and Baka (B) cohorts, both of which exhibit variation in the DARC genotype. The red line represents the linear regression line. We report the Pearson correlation coefficient and significance of that correlation. We observe a slightly negative but non-significant relationship between neutrophil proportion and Duffy null genotype count.

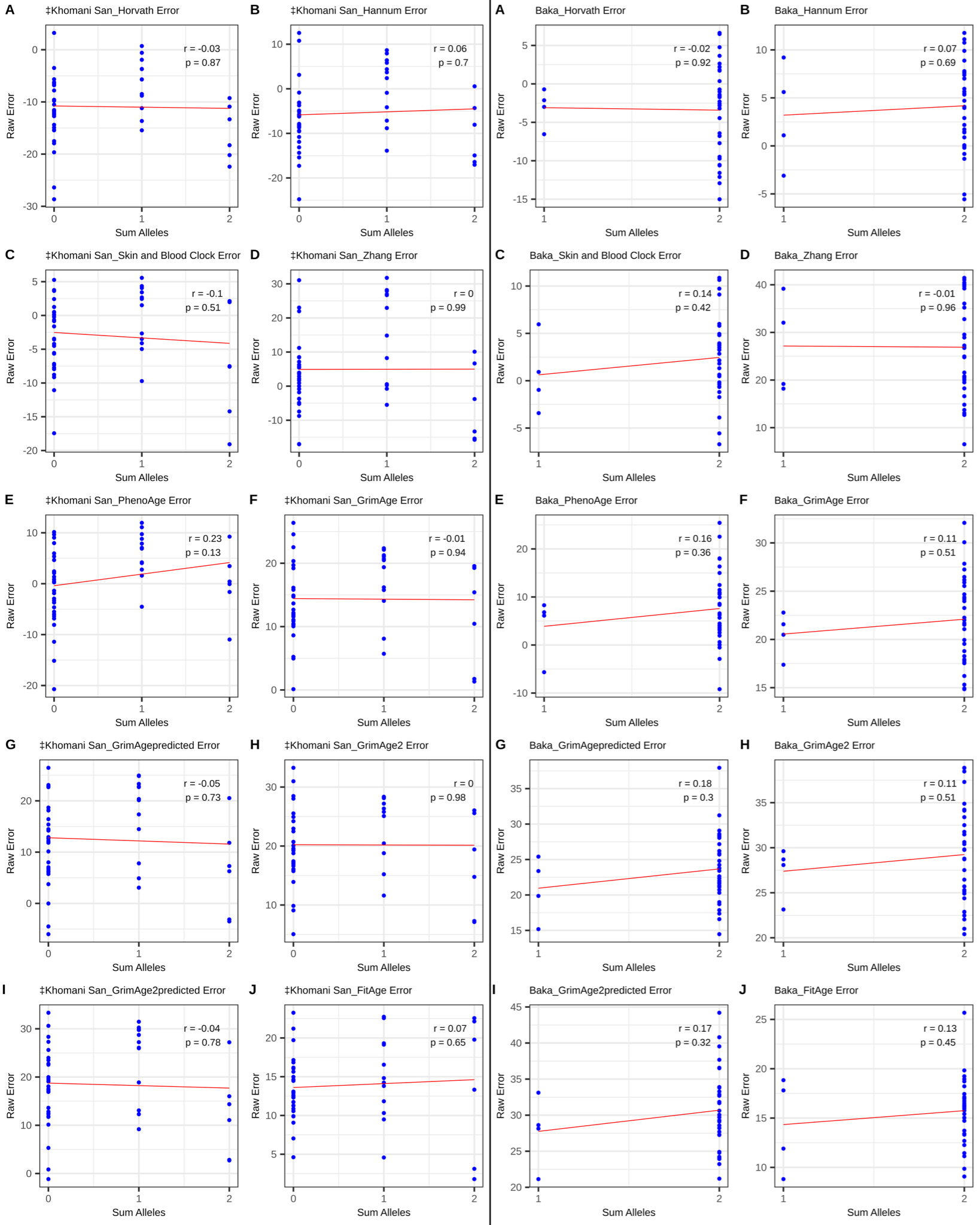

#### Supplementary Figure 4 - Correlation of DARC genotype status with prediction error across epigenetic clocks

We find that there is no systematic relationship between Duffy null count, plotted on the x-axis, and prediction error, plotted on the y-axis, from 10 age prediction methods across the ‡Khomani San (left) and Baka (right) cohorts. The red line represents the linear regression line. We report the Pearson correlation coefficient and significance of that correlation.

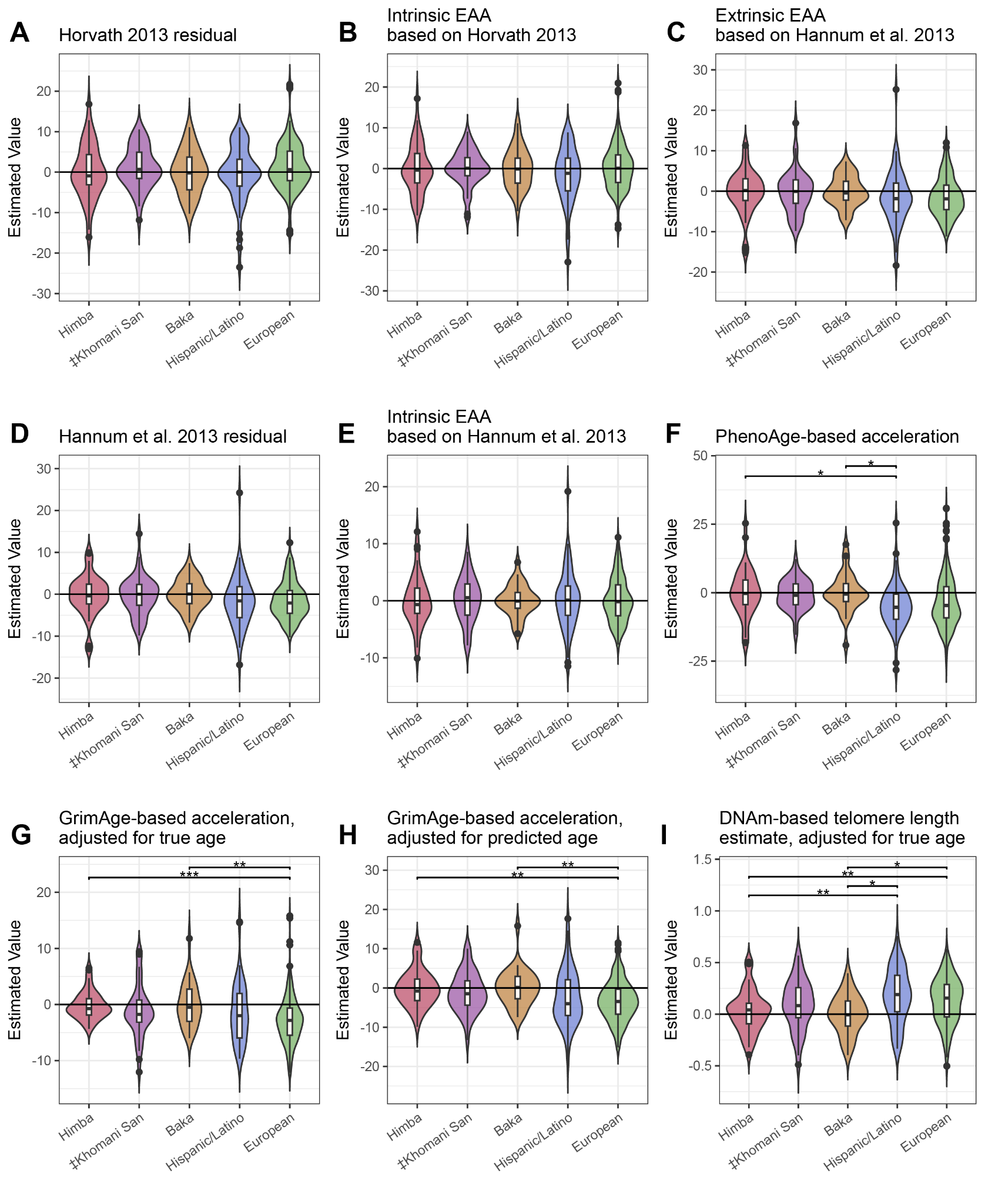

#### Supplementary Figure 5 - Distributions of epigenetic age acceleration estimates across diverse cohorts

Violin plots A-H show the distributions of epigenetic age acceleration estimates from 8 different methods among Himba, ‡Khomani San, Baka, European, and Hispanic cohorts . Panel I shows a methylation-based estimate of telomere length after adjusting for chronological age. A higher value here indicates longer estimated telomere length than expected based on chronological age. We tested for significant differences in epigenetic age acceleration among all populations by ANOVA, followed by a Tukey test to identify significant pairwise differences. * indicates an adjusted p-value of < .05, ** < .01, and *** < .001.

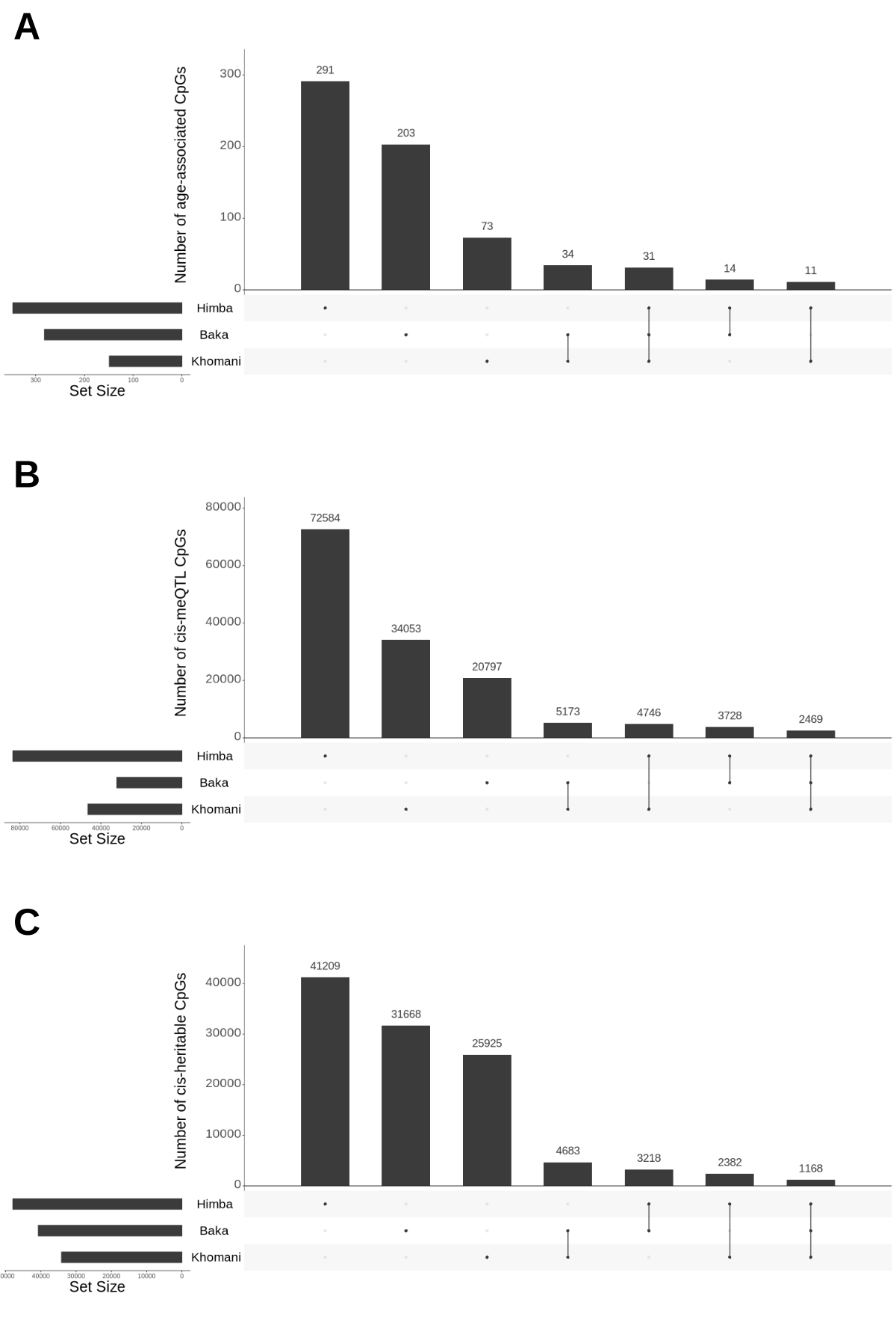

#### Supplementary Figure 6 - Overlap of significant CpG sites across the Himba, ‡Khomani San, Baka

Upset plots showing the overlap of significant A) age-associated CpGs from the unadjusted EWAS B) *cis*-meQTL influenced CpGs from the baseline scan and c) *cis*-heritable CpGs from the FUSION-based meQTL scan among the 3 populations.

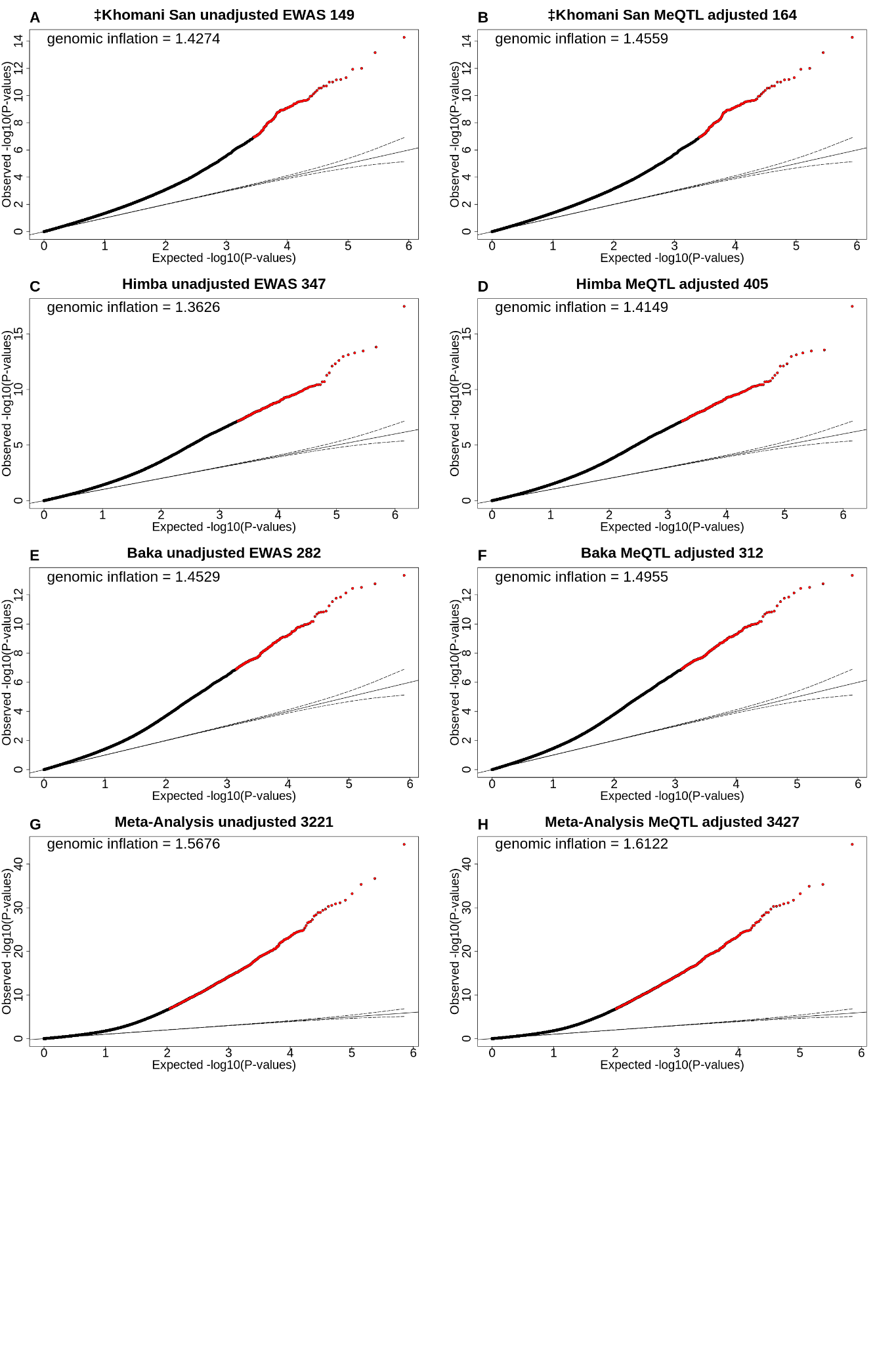

#### Supplementary Figure 7 - Genomic Inflation comparisons for meQTL-adjusted vs unadjusted EWAS

Quantile-quantile plots for each population-specific EWAS and the meta-analysis. Expected p-values under a uniform distribution are plotted on the x-axis, while the observed p-values are plotted on the y-axis. The left column shows the results of the unadjusted EWAS and the right column shows the results of the top meQTL-adjusted EWAS. The number of significantly associated CpG sites is listed in each plot title.

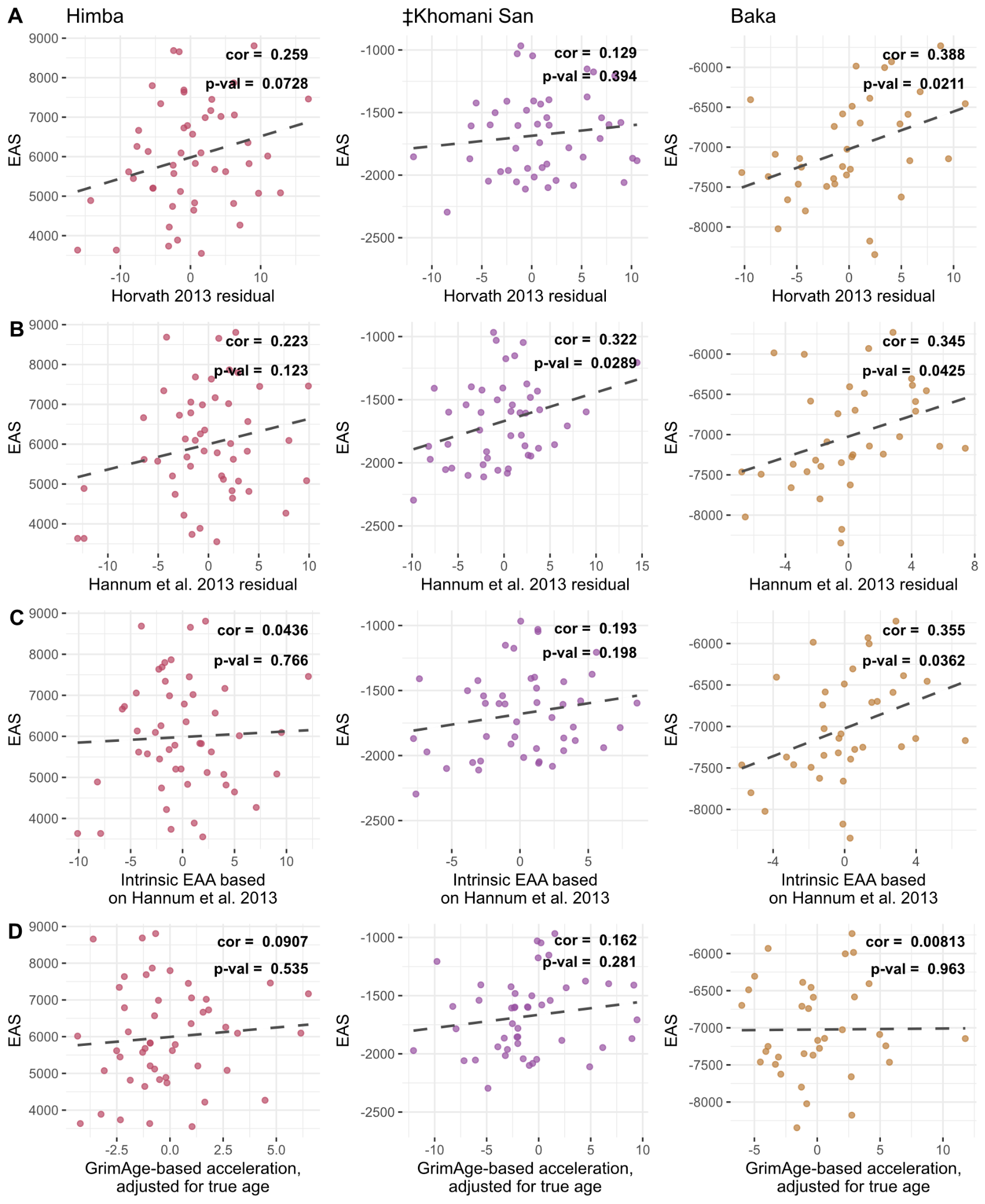

#### Supplementary Figure 8 - Relationship between EAS and metrics of epigenetic age acceleration

Scatterplots show the relationship between the genotype-based epigenetic aging score (EAS) and 4 additional metrics of age for the Himba, ≠Khomani San, and Baka (see Figure 7). Individuals’ EAS values were plotted against A) the Horvath 2013 clock and B) Hannum et al. 2013 clock residuals after correcting for chronological age, C) ‘Intrinsic Epigenetic Age Acceleration’, based on the Hannum et al. 2013 clock, and D) epigenetic age acceleration based on GrimAge adjusted for true age.

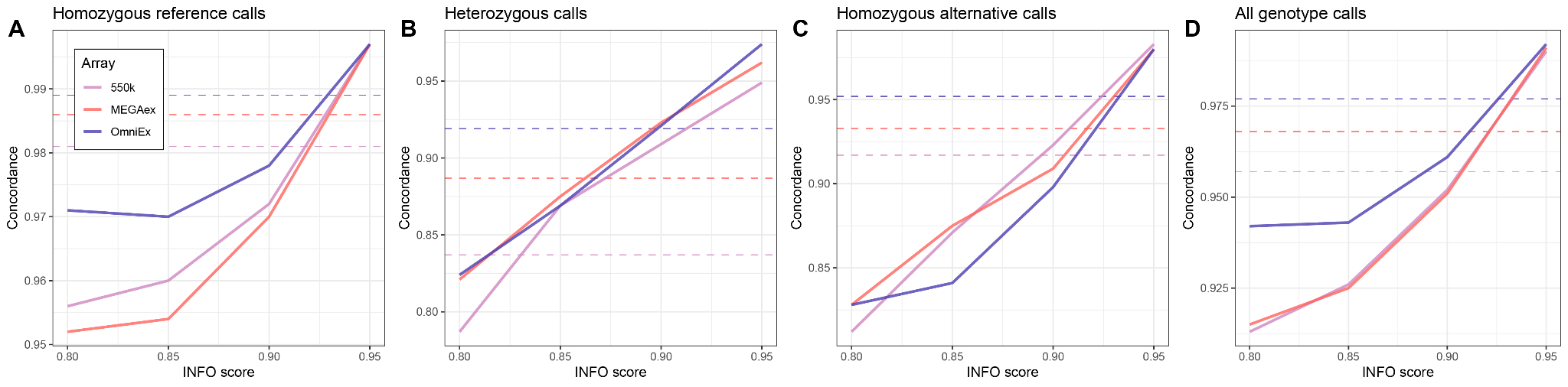

#### Supplementary Figure 9 - Concordance of imputed genotypes with exome calls by INFO score.

The proportion of concordant genotype calls between imputed data and independently generated exome data is plotted along the y-axis for 37 ‡Khomani San individuals, binned by imputation INFO score, plotted along the x-axis. The INFO score generally reflects imputation quality. The solid lines represent the average concordance values for individuals genotyped on each array. The dashed horizontal lines correspond to the average concordance across all INFO scores.

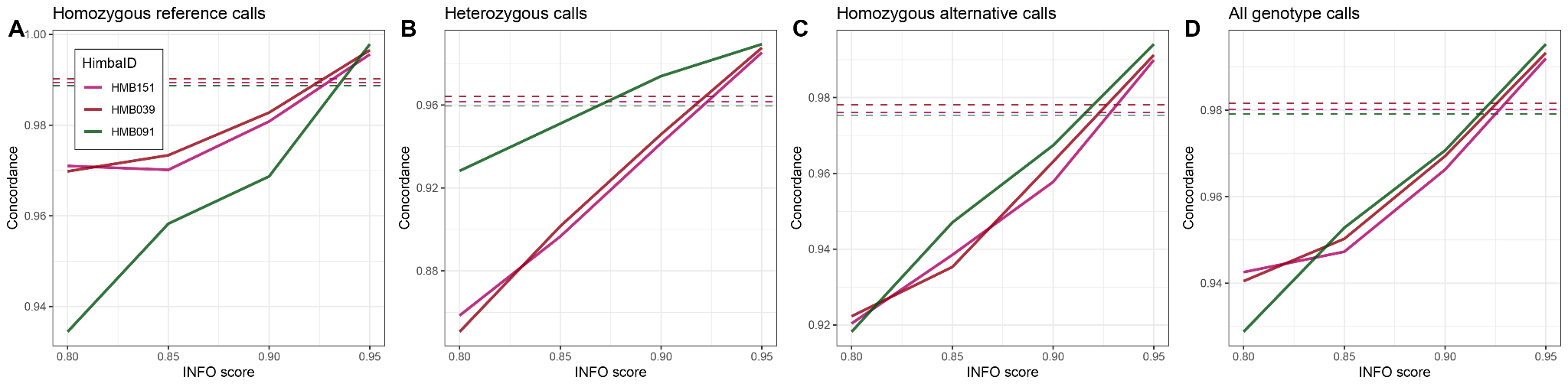

#### Supplementary Figure 10 - Concordance of imputed genotypes with independent calls from genotyping on the MEGAex array by INFO score

The proportion of concordant genotype calls between imputed data and independent genotyping array data is plotted along the y-axis, binned by imputation INFO score, plotted along the x-axis. The INFO score generally reflects imputation quality. The solid lines represent the concordance values for 3 Himba individuals that were typed on both the H3Africa and the MEGAex arrays. The dashed horizontal lines correspond to the average concordance across all INFO scores.

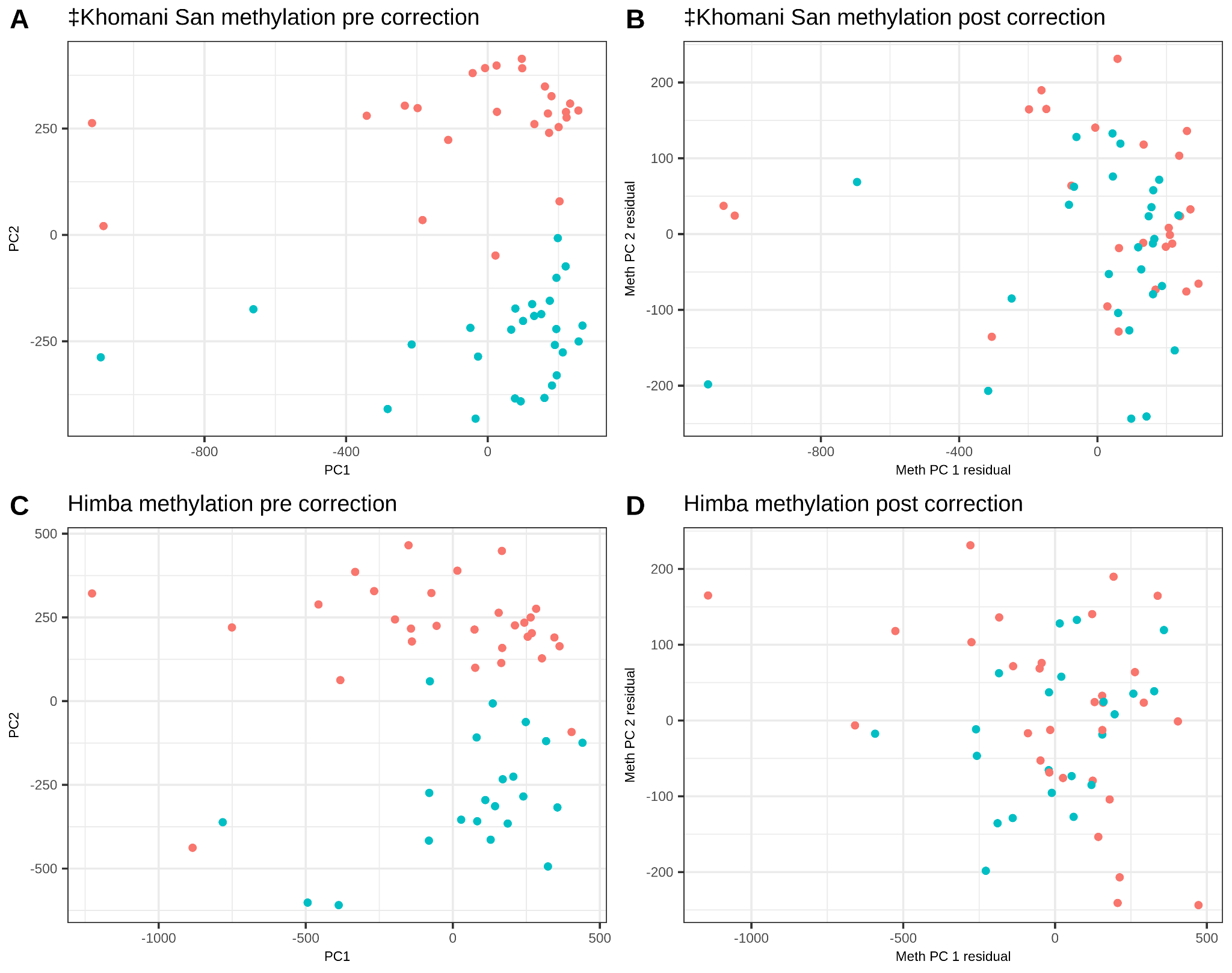

#### Supplementary Figure 11 - Principal component analysis of batch effects in DNA methylation data

The first and second principal components are plotted on the x- and y-axis, respectively. Batch effects drive clustering in the second principal component in both the ‡Khomani San (A) and Himba © DNA methylation data. Red and blue points represent the two batches for each cohort. Regressing out batch number (B) and the first 20 control probe intensity PCs (D) from the DNA methylation PCs of the ‡Khomani San and the Himba, respectively, reduces clustering by batch.

#### Supplementary Figure 12 - Principal components analysis of genotype data

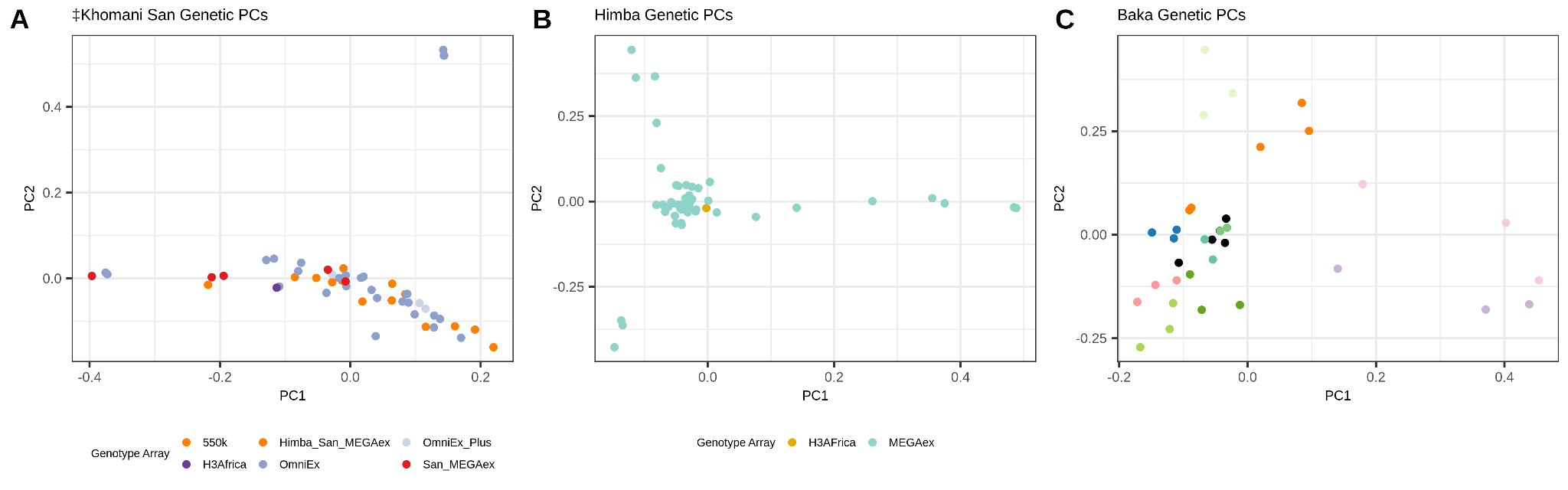

The first and second principal components are plotted on the x- and y-axis, respectively. We observe no clustering by genotype array in the ‡Khomani San and Himba. The Baka data show strong evidence of clustering according to parent-offspring or trio membership, indicated by clusters of coloured dots (black dots represent unrelated individuals).

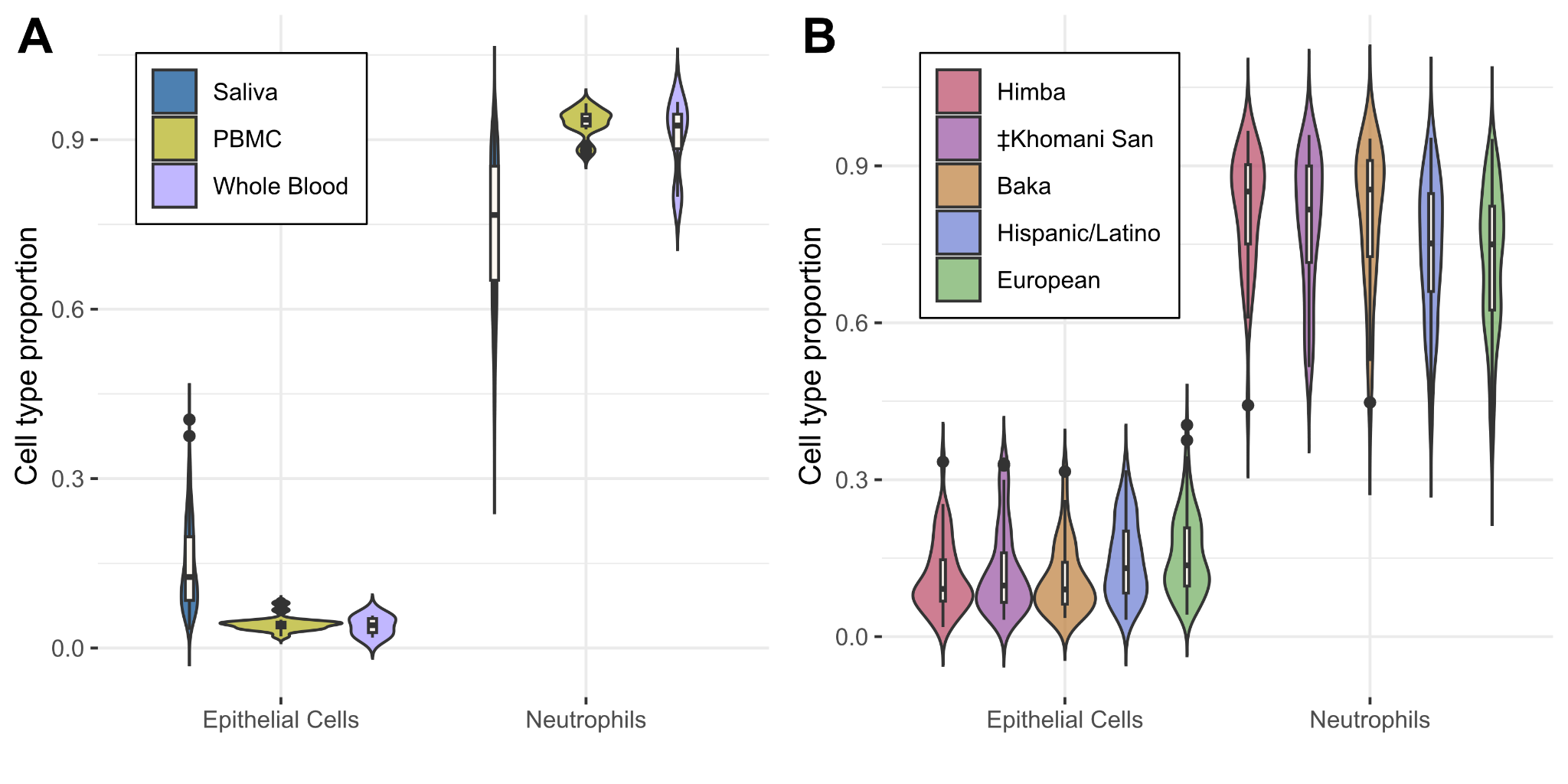

#### Supplementary Figure 13 - Estimated cell type composition by tissue type and population

The estimated proportions of epithelial and neutrophil cells , the two most abundant cell types in saliva, are plotted on the y-axis, grouped by either the estimated tissue type (A) or population (B). A) Samples predicted to be either PBMC (n = 17) or whole blood (n = 4) have cell type proportions that fall within the range of samples predicted to be saliva (n = 308). B) Cell type proportions do not vary systematically by population for samples predicted to be derived from either saliva, PBMCs, or whole blood.

#### Supplementary Figure 14 - Schematic of the epigenetic aging score

A cartoon depiction of how the direction of the effect of the SNP on DNA methylation and the direction of effect of DNA methylation on age combine to produce a consistent overall effect that is summed in the epigenetic aging score.

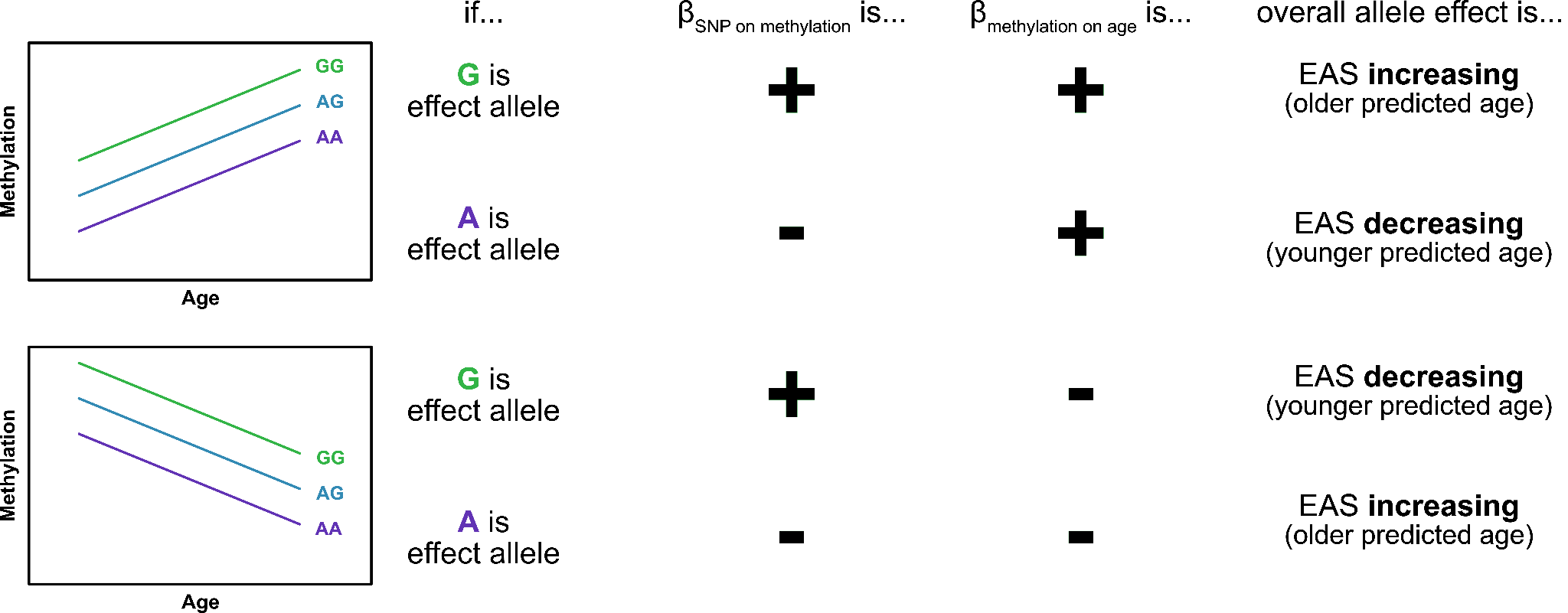

### Supplementary Tables

#### Supplementary Table 1

Mean age-adjusted prediction error from published 10 epigenetic age estimation methods for the Himba, ‡Khomani San, Baka, European, and Hispanic/Latino cohorts (Figure 1).

|  | **Himba** | **‡Khomani San** | **Baka** | **Hispanic** | **European** |
| --- | --- | --- | --- | --- | --- |
| **Horvath** | -0.34 | -1.1 | 0 | -1.15 | 1.13 |
| **Hannum et. al 2013** | -4.55 | -1.99 | 0.43 | 1.34 | 1.59 |
| **Skin and Blood Clock** | -6.53 | -0.68 | 1.74 | 1.09 | 1.65 |
| **Zhang et. al 2019** | -2.36 | -1.09 | 0.35 | 0.54 | 0.9 |
| **PhenoAge** | -10.23 | -0.16 | 4.14 | 0.89 | 2.33 |
| **GrimAge** | -0.73 | 7.55 | 4.84 | -1.41 | -2.95 |
| **GrimAgepredicted** | -4.94 | 7.11 | 5.97 | -0.71 | -1.89 |
| **GrimAge2** | 0.5 | 6.94 | 3.92 | -1.17 | -3.08 |
| **GrimAge2predicted** | -3.34 | 6.54 | 4.95 | -0.53 | -2.1 |
| **FitAge** | -1.6 | 4.33 | 2.42 | -0.87 | -1.14 |

#### Supplementary Table 2

Mean raw unadjusted prediction error (mean absolute prediction error) from published 10 epigenetic age estimation methods for the Himba, ‡Khomani San, Baka, European, and Hispanic/Latino cohorts (Supplementary Figure 2).

|  | **Himba** | **‡Khomani San** | **Baka** | **European** | **HispanicLatino** |
| --- | --- | --- | --- | --- | --- |
| **Horvath** | -7.37 (9.22) | -10.9 (11.07) | -3.38 (5.11) | -10.7 (11.08) | -12.99 (12.99) |
| **Hannum** | -4.97 (8.87) | -5.49 (8.37) | 4.07 (4.99) | -4.17 (5.43) | -4.44 (6.13) |
| **Skin and Blood Clock** | -7.6 (7.96) | -2.94 (4.93) | 2.25 (3.69) | -1.48 (3.42) | -2.05 (4.21) |
| **Zhang** | 12.53 (18.08) | 4.93 (10.42) | 26.92 (26.92) | 0.44 (5.11) | -0.03 (6.68) |
| **PhenoAge** | -8.38 (9.95) | 0.79 (5.8) | 7.17 (8.21) | 2.64 (7.19) | 1.18 (6.08) |
| **GrimAge on true age** | 10.51 (10.8) | 14.37 (14.37) | 21.92 (21.92) | 0.64 (3.3) | 2.15 (4.44) |
| **GrimAge on predicted age** | 5.62 (10.32) | 12.48 (13.23) | 23.36 (23.36) | -0.32 (4.36) | 0.83 (5.86) |
| **GrimAge2 on true age** | 18.87 (18.87) | 20.2 (20.2) | 29.03 (29.03) | 6.45 (6.92) | 8.32 (8.88) |
| **GrimAge2 on predicted age** | 14.4 (15.45) | 18.47 (18.52) | 30.35 (30.35) | 5.58 (6.79) | 7.11 (8.88) |
| **FitAge** | 9.51 (9.71) | 13.87 (13.87) | 15.59 (15.59) | 7.26 (7.45) | 7.5 (7.97) |

#### Supplementary Table 3

A list of previously published studies DNA methylation and age indicating the focal tissue type and total number of associations.

| **Study** | **Tissue** | **Number of associations** |
| --- | --- | --- |
| Alisch et al., 2012 | Peripheral blood | 2078 |
| Bell et al., 2012 | Whole Blood | 490 |
| Bocklandt et al., 2011 | Saliva | 88 |
| Cruickshank et al., 2013 | Whole Blood | 3244 |
| Fernández et al., 2015 | Mesenchymal Stem Cells | 60155 |
| Florath et al., 2013 | Whole Blood | 162 |
| Garagnani et al., 2012 | Whole Blood | 9 |
| Hannum et al., 2013 | Whole Blood | 71 |
| Heyn et al., 2012 | Whole Blood | 3205 |
| Johansson et al., 2013 | Leukocytes | 137658 |
| Kananen et al., 2016 | Whole Blood | 1202 |
| Marttila et al., 2015 | Leukocytes | 8540 |
| Raykan et al., 2010 | Whole Blood | 131 |
| Smith et al., 2014 | Peripheral Blood | 30 |
| Teschendorff et al., 2010 | Whole Blood | 589 |
| Weidner et al., 2014 | Whole Blood | 102 |
| Xu et al., 2014 | Whole Blood | 749 |
| Zaghool et al., 2015 | Whole Blood | 828 |
| Horvath 2013 | Multi-tissue | 353 |
| Han et al., 2020 | Whole Blood | 65 |
| Lee et al., 2020 | Whole Blood | 4367 |
| Koch et al., 2011 | Dermis, epidermis, cervical smear, T-cells, monocytes | 20 |
| Zhang et al., 2019 | Whole blood, saliva | 514 |
| Bekaert et al., 2015 | Whole blood, teeth | 7 |
| Florath et al., 2014 | Whole blood | 162 |
| Li et al., 2018 (a) | Whole blood | 6349 |
| Li et al., 2018 (b) | Whole blood | 6 |
| Li et al., 2018 (c) | Whole Blood | 238 |
| Naue et al., 2017 | Whole Blood | 9 |
| Vidal-Bralo et al., 2016 | Whole Blood | 8 |
| Weidner et al., 2014 | Whole Blood | 3 |
| Wu et al., 2019 | Whole Blood | 111 |
| Xu et al., 2015 | Whole Blood | 2965 |
| Horvath et al., 2018 | Skin and Whole Blood | 391 |
| Gopalan et al. 2017 | Saliva and Whole blood | 107 (novel) |

#### Supplementary Table 4

A table of counts for the number of times a particular regression method produced the best fitting meQTL model in FUSION analysis, determined by the highest mean R-squared after 5-fold cross-validation.

| **Population** | **Elastic Net** | **SuSie** | **LASSO** | **Top1** |
| --- | --- | --- | --- | --- |
| ‡Khomani | 27187 | 3486 | 1428 | 2057 |
| Baka | 38926 | 1012 | 344 | 455 |
| Himba | 47912 | 50 | 0 | 15 |
| Combined | 20169 | 10368 | 4457 | 6442 |

#### Supplementary Table 5

Raw error of the age prediction models trained on CpG sites without significant meQTL or *cis*-heritability (non-heritable) and those with significant meQTL or *cis*-heritability (heritable predictors), averaged over 100 models each. Standard deviation shown in parentheses. P-values were determined by a two-sided T-test comparing the non-heritable and heritable distributions of average error.

|  | **Non-heritable predictors** | **Heritable predictors** | **P-value** |
| --- | --- | --- | --- |
| **African test samples** | 0.02 years (0.92) | 0.27 years (1.03) | 0.066 |
| **European samples** | -1.83 years (2.58) | 3.19 years (3.11) | <2.2e-16 |
| **Hispanic/Latino samples** | -2.88 years (2.73) | 2.4 years (3.29) | 2.7e-02 |

#### Supplementary Table 6

A table of the number of individuals from each cohort in each DNA methylation typing batch.

| **Batch** | **Methylation Array** | **Population sample size** |
| --- | --- | --- |
| 1 | Illumina 450k | Baka (n = 36), ‡Khomani San (n = 28) |
| 2 | Illumina 450k | ‡Khomani San (n = 29) |
| 3 | Illumina EPIC | Himba (n = 30) |
| 4 | Illumina EPIC | Himba (n = 21) |

#### Supplementary Table 7

A summary table of DNA methylation array quality control metrics by batch

|  | **Batch 1 (‡Khomani San and Baka)** | **Batch 2 (‡Khomani San)** | **Batch 3**  **(Himba)** | **Batch 4**  **(Himba)** |
| --- | --- | --- | --- | --- |
| **SNP probes** | 65 | 65 | 59 | 59 |
| **Non-autosomal probes** | 11656 | 11656 | 19646 | 19646 |
| **Probes failing in >5% of samples** | 1606 | 1201 | 2061 | 2544 |
| **Pooly mapping probes** | 40490 | 40490 | 63483 | 63483 |
| **Non-unique 30mer probes** | 16546 | 16546 | 23428 | 23428 |
| **Non-CpG probes** | 3091 | 3091 | 2932 | 2932 |

#### Supplementary Table 8

A table of the number of individuals from each cohort typed on each genotyping array.

| **Populations** | **Genotype Array** | **# Samples used** | **Number of variants pre-imputation** |
| --- | --- | --- | --- |
| ‡Khomani San | Illumina 550Kv2 | 12 | 483,740 |
| ‡Khomani San | OmniExpress | 29 | 671,097 |
| ‡Khomani San | OmniExPlus | 3 | 667,221 |
| ‡Khomani San | MEGAex | 5 | 717,947 |
| ‡Khomani San | H3Africa | 1 | 1,480,928 |
| Himba | MEGAex | 49 | 748,985 |
| Himba | H3Africa | 1 | 1,644,188 |
| ‡Khomani San, Himba | MEGAex | 2, 1 | 716,989 |
| Baka | OmniOne | 35 | 721,083 |

#### Supplementary Table 9

Reference-based cell type deconvolution for replicate samples. ‡Khomani and Himba replicates were typed in different batches. Himba 181r was sampled again 3 years after the original collection.

|  | **Epithelial** | **Fibroblast** | **NK** | **B** | **CD4T** | **CD8T** | **Monocytes** | **Neutrophils** |
| --- | --- | --- | --- | --- | --- | --- | --- | --- |
| Baka 03F | .124 | .026 | .015 | 0 | .005 | 0 | .018 | .812 |
| Baka 03Fr | .12 | .023 | .017 | 0 | .01 | 0 | .019 | .81 |
| Baka 13F | .036 | .012 | 0 | 0 | 0 | 0 | 0 | .952 |
| Baka 13Fr | .035 | .014 | 0 | 0 | 0 | 0 | 0 | .951 |
| Khomani 4 | .168 | .028 | .053 | .037 | .033 | 0 | .094 | .587 |
| Khomani 4r | .175 | .028 | .06 | .043 | .044 | 0 | .1 | .55 |
| Himba 181 | .017 | .011 | .005 | 0 | 0 | 0 | 0 | .967 |
| Himba 181r | .02 | .013 | .003 | 0 | 0 | 0 | 0 | .964 |
